# Do initial concentration and activated sludge seasonality affect pharmaceutical biodegradation rate constants?

**DOI:** 10.1101/2020.12.17.423224

**Authors:** Tamara J.H.M. van Bergen, Ana B. Rios-Miguel, Tom M. Nolte, Ad M.J. Ragas, Rosalie van Zelm, Martien Graumans, Paul Scheepers, Mike S.M. Jetten, A. Jan Hendriks, Cornelia U. Welte

## Abstract

Pharmaceuticals find their way to the aquatic environment via wastewater treatment plants (WWTPs) and biodegradation plays an important role in mitigating environmental risks, however a mechanistic understanding of involved processes is limited. The aim of this study was to evaluate potential relationships between first-order biodegradation rate constants (k_b_) of nine pharmaceuticals and initial concentration of the selected compounds, and sampling season of the used activated sludge inocula. Four-day bottle experiments were performed with activated sludge from WWTP Groesbeek (The Netherlands) of two different seasons, summer and winter, spiked with two environmentally relevant concentrations (3 and 30 nM) of pharmaceuticals. Concentrations of the compounds were measured by LC-MS/MS, microbial community composition was assessed by 16S rRNA gene amplicon sequencing and k_b_ values were calculated. The biodegradable pharmaceuticals, ranked from high to low biodegradation rates, were acetaminophen, metformin, metoprolol, terbutaline, and phenazone. Carbamazepine, diatrizoic acid, diclofenac and fluoxetine were not converted. Summer and winter inocula did not show significant differences in microbial community composition, but resulted in a slightly different k_b_ for some pharmaceuticals. Likely microbial activity was responsible instead of community composition. In the same inoculum different k_b_ values were measured, depending on initial concentration. In general, biodegradable compounds had a higher k_**b**_ when the initial concentration was higher. This demonstrates that Michealis-Menten kinetics theory has shortcomings for some pharmaceuticals at low, environmentally relevant concentrations and that the pharmaceutical concentration should be taken into account when measuring the k_b_ in order to reliably predict the fate of pharmaceuticals in the WWTP.

## 1. Introduction

Tonnes of pharmaceuticals are prescribed annually, which partly find their way to the aquatic environment via sewage systems and wastewater treatment plants (WWTPs) (Fent et al. 2006). The concentration of these pharmaceuticals in the aquatic environment is generally low (ranging from nM to μM concentrations), and thus they are considered as organic micropollutants (OMPs). Even at such low concentrations, they can elicit large adverse effects on aquatic life including long-term or short-term toxicity (e.g. Fent et al. 2006, Santos et al. 2010). In WWTPs, the fate of a pharmaceutical is mostly influenced by microbial biodegradation (Verlicchi et al. 2012), a process which is still difficult to capture accurately with modelling approaches. Statistical quantitative structure activity relationship (QSAR) models can describe approximately 50% of the variability in biodegradation rate constants (e.g. Nolte et al. 2020). This explains for a large part why some OMPs, including pharmaceuticals, are better removed than others. However, the question remains why some WWTPs remove specific pharmaceuticals better than others. Statistical meta-analyses either find a low explanatory value for the removal efficiency of a large structurally diverse set of OMPs, including pharmaceuticals (17%; Douziech et al. 2018), or perform better but are limited to specific groups of OMPs, including (R2adj ranged from 0.35-0.73; Wang et al. 2020). Due to the limited knowledge on general principles influencing microbial conversion of pharmaceuticals, biodegradation is yet not modelled mechanistically in fate models such as SimpleTreat, which is part of the European Union System for the Evaluation of Substances (Franco et al. 2013, Struijs 2014).

Instead, biodegradation rate constants used in fate models are typically estimated with standard methods such as OECD tests (OECD TG 301 series, TG 310, TG 302 series, TG 314B and TG 303A, in case of SimpleTreat). It has to be noted that these tests are not designed for this purpose, but rate constants can nevertheless be derived from percentage removal or biodegradability categories (Struijs 2014). When OECD test outcomes are used to predict the fate in WWTPs, the environmental realism of the biodegradation rate constants is questionable as the test outcomes apply to specific incubation conditions with a long time duration and the microbial community composition of the inoculum varies (Goss et al. 2020, Kowalczyk et al. 2015, Li and McLachlan 2019, Rücker and Kümmerer 2012). In most OECD screening tests pharmaceutical concentrations of 10–400 mg of carbon per liter may be applied (Kowalczyk et al. 2015), which is far above environmentally relevant concentrations (in the range of μg L^−1^) and this may affect biodegradation rate constants. Recent studies showed that first-order biodegradation rate constants depend on initial concentration in activated sludge treatment plants (Nolte et al. 2020) and in biofilm reactors (Svendsen et al. 2020). Biodegradation rate constants are often calculated as pseudo-first order constants (hereafter referred to as kb, Schwarzenbach et al. 2005), assuming no saturation of enzymes occurs and the maximum velocity of the reaction is not reached (for more information, see 2.4.1). In the calculation of k_b_, an effect of concentration is already taken into account as the decrease in concentration over time depends on a rate constant and the concentration. Svendsen et al. (2020) showed that for some pharmaceuticals, k_b_ did not follow typical Michaelis-Menten kinetics at low, environmentally relevant concentrations (1-10 μg L^−1^) as k_b_ is expected to stay constant and the removal rate decreases over time as a result of the changing concentration. Instead, the k_**b**_ of some pharmaceuticals such as metoprolol increased with increasing initial concentration.

The concentration of pharmaceutical compounds in the influent of WWTPs can vary between seasons as a result of different consumption patterns by the population and a different amount of rainfall (Caucci et al. 2016, Di Marcantonio et al. 2020). Furthermore, WWTP operational parameters (i.e. solid retention time and hydraulic retention time) and other environmental conditions such as temperature have seasonal variability (Awolusi et al. 2018, Limpiyakorn et al. 2005). As a consequence of these changes, different microbial community compositions have been found in same WWTPs at different seasons. For example, ammonia oxidizing bacteria (AOB) were often found more abundant during summer than during winter and this was correlated to higher nitrification rates and lower ammonium concentration in the effluent of WWTPs (Awolusi et al. 2018, Ju et al. 2014, Liu et al. 2019, Wang et al. 2012). Previous experiments have observed a positive correlation between the k_b_ of specific pharmaceuticals and ammonium removal or nitrification rate (Fernandez-Fontaina et al. 2012, Helbling et al. 2012). Furthermore, higher abundance of specific microbial taxa has been correlated to a higher k_b_ and removal efficiency of some pharmaceuticals in AS (Helbling et al. 2015, Kim et al. 2017).

The aim of this study was to determine the influence of initial pharmaceutical concentration and sampling season of AS inocula on the k_b_ of pharmaceutical compounds. To our knowledge, this is the first time that the effect of pharmaceutical concentration on k_b_ is experimentally studied in activated sludge by means of a batch test. We selected nine pharmaceuticals based on high prescription rates in Europe (Fent et al. 2006) and/or with a high priority to monitor due to adverse environmental effects (i.e. pharmaceuticals on EU and Dutch watch lists): acetaminophen, metformin, diclofenac, metoprolol, terbutaline, diatrizoic acid, fluoxetine, and carbamazepine. Four-day batch assays were performed with AS inocula of the same WWTP taken at different seasons (summer and winter). Biodegradation rate constants and solids water partition coefficients (k_b_ and k_d_, respectively) were generated at two environmentally relevant concentrations (3 and 30 nM). Summer and winter samples were analyzed on nitrification activity and microbial community. We hypothesized that i) a higher spiked concentration in the range of 3 to 30 nM result in a higher k_b_; ii) inocula taken at different seasons in the same WWTP have different characteristics (i.e. microbial community, nitrification activity and pharmaceutical concentration) that will affect the biodegradation rate constants of pharmaceuticals, with higher rate constants during summer.

## 2. Materials and methods

### 2.1. Chemical selection and experimental setup

We conducted biodegradation assays with nine pharmaceuticals (Table 1) twice: in June 2019 (summer) and December 2019 (winter). The log K_ow_ was obtained from Drugbank (Wishart et al. 2006), while information on the solid form of the pharmaceuticals was provided by the supplier (SI 1).

**Table 1.** Information on the pharmaceuticals selected in the experiment and the solid form that was used to obtain the dissolved spiking mixture. Log K_ow_ is the octanol water partition coefficient.

| Pharmaceutical | Cas number | Class | Log $K_{ow}$ | Solid form used for experiment | CAS number solid form | Molecular mass solid form (g/mol) | Supplier's name | Product number | Purity |
| --- | --- | --- | --- | --- | --- | --- | --- | --- | --- |
| Acetaminophen | 103-90-2 | Acid | 0.46 | Acetaminophen | 103-90-2 | 151.2 | Sigma Aldrich | A7085 | ≥99% |
| Carbamazepine | 298-46-4 | Neutral | 2.45 | Carbamazepine | 298-46-4 | 236.3 | Sigma Aldrich | C4024 | 100% |
| Diatrizoic acid | 117-96-4 | Acid | 1.37 | Diatrizoic acid | 117-96-4 | 613.9 | Sigma Aldrich | D9268 | 100% |
| Diclofenac | 15307-86-5 | Acid | 1.85 | Diclofenac sodium salt | 15307-79-6 | 318.1 | Sigma Aldrich | D6899 | 100% |
| Fluoxetine | 54910-89-3 | Base | 1.51 | Fluoxetine Hydrochloride | 56296-78-7 | 345.8 | Sigma Aldrich | PHR13 94 | 100% |
| Metformin | 657-24-9 | Base | -4.13 | 1,1 dimethylbiguanide hydrochloride | 1115-70-4 | 165.6 | Sigma Aldrich | D5035 | ≥ 97% |
| Metoprolol | 37350-58-6 | Base | 1.88 | Metoprolol Tartrate salt | 56392-17-7 | 684.8 | Sigma Aldrich | 80337 | 100% |
| Phenazone | 60-80-0 | Base | 0.38 | Phenazone | 60-80-0 | 188.2 | Sigma Aldrich | A5882 | 100% |
| Terbutaline | 23031-25-6 | Base | 0.9 | Terbutaline Sulfate | 23031-32-5 | 274.3 | Sigma Aldrich | T00500 00 | 100% |

The removal of pharmaceuticals was tested for a duration of 96 h, which is sufficient to observe short term activity due to the high microbial activity of undiluted activated sludge samples (e.g. Helbling et al. 2010). As the AS samples were undiluted and microorganisms were not washed in order to obtain environmental realism of the test, no lag phase was anticipated on. We tested three treatments in summer and winter AS inocula (Figure 1):

1. “AS” treatment: where we evaluated pharmaceutical biodegradation at low concentrations in AS. In the summer experiment we measured the removal of background concentrations and in the winter experiment we spiked 3nM of each pharmaceutical.
2. “AS30” treatment: where we evaluated pharmaceutical biodegradation a high concentration in AS. To do that, we spiked with 30 nM of each pharmaceutical in both summer and winter experiments.
3. “iAS30” treatment: where we assessed the abiotic removal of pharmaceuticals in AS (i.e. sorption). For that we doubled autoclaved AS and spiked 30 nM of each pharmaceutical.

**Figure 1.**
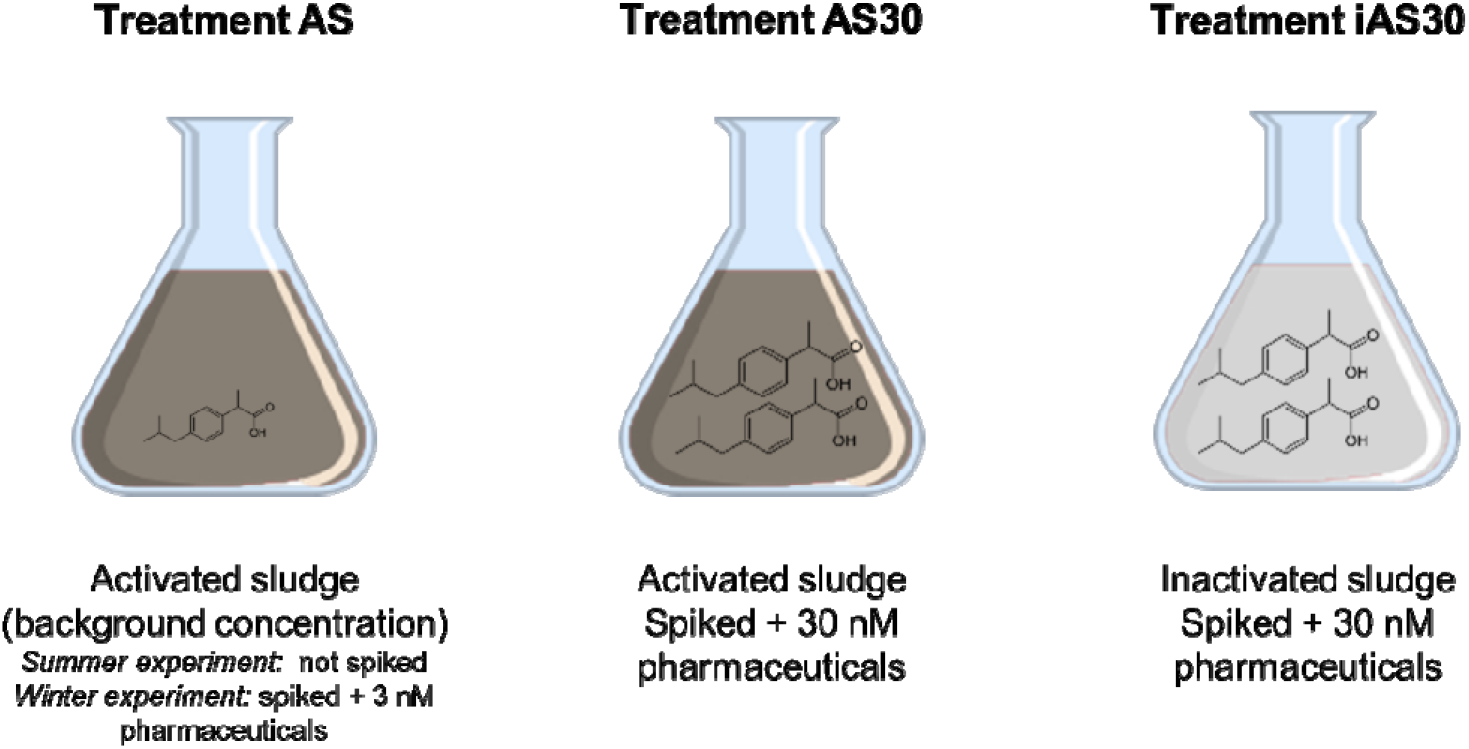
Treatments used in the summer and winter experiments: AS was either non-spiked, only including the background concentration of pharmaceuticals (Summer experiment), or spiked with 3 nM pharmaceuticals (Winter experiment); AS30 is the AS treatment that was spiked with 30 nM of pharmaceuticals; and iAS30 is the inactivated sludge treatment that was spiked with 30 nM of pharmaceuticals.

The biotransformation experiments were carried out in 120 mL glass bottles. AS samples were taken from WWTP Groesbeek (Gelderland, The Netherlands), identified as a national hotspot of OMPs (Vissers et al. 2017). This WWTP has a maximum hydraulic capacity of 900 m^3^/h and it treats mostly domestic water (estimated 90%). The WWTP contains an aeration tanks with a sludge age of 12 days. Five liters of activated sludge were sampled in Winter and Summer time less than 72 hours before the start of the experiment. The sample was immediately transported to the laboratory where it was stored at 4 °C till the start of the experiment (t = 0). Individual stock solutions were prepared in methanol for each pharmaceutical at a concentration of 3mM. Afterwards, a mixture of the nine pharmaceuticals was prepared in methanol too (final concentration of each pharmaceutical: 0.3 mM). To minimize the potential effects of the methanol solvent, the spike mixture of pharmaceuticals (to a final bottle concentration of 3 or 30 nM) was added to each bottle prior to addition of sampled AS. Methanol was allowed to evaporate in a fume hood with gentle air circulation for approximately 30 minutes and subsequently 90 mL of AS or iAS were added to the bottles. Additionally, 50 mM HEPES (4-(2-hydroxyethyl)-1-piperazineethanesulfonic acid) was added in order to prevent acidification and maintain a pH around 7. Prior incubations showed a drop in pH below 6 leading to inhibition of nitrification (Ruiz et al. 2003) with possible consequences for biodegradation rates. The bottles were manually mixed to allow for complete dissolution of pharmaceuticals and the first samples were taken (t = 0) and centrifuged to separate the sludge from the aqueous phase. Hereafter, samples were taken after 4, 8, 24, 48 and 96 h. Bottles were closed with cotton plugs to allow aeration and prevent contamination from the environment. The bottles were incubated at 15 °C and shaken at 200 RPM to ensure continuous mixing and aeration, so that a constant dissolved oxygen concentration of approximately 1 mg L^−1^ was maintained. All biotransformation assays were run in triplicates. Bottles were incubated in the dark in order to reflect WWTP conditions and to exclude photodegradation or growth of photoautotrophs. Double autoclaved inactivated sludge bottles (treatment iAS30) were prepared before the start of the experiment (and spiking) by autoclaving bottles at 121 °C and 103 kPa for 20 min and repeating this procedure after 24 h as previously described (Helbling et al. 2010).

### 2.2. Wastewater analyses of the batch incubation

Temperature and pH were monitored every 24h with a Metrohm Applikon pH meter (Schiedam, the Netherlands). In the beginning and at the end of the experiment (t = 0 and 96h), the dissolved organic and inorganic carbon concentrations (DOC and DIC, respectively) were measured, as well as total suspended solids. Water samples for DIC analyses were filtered with glass-fiber filters (Ø 0.45 μm), stored at 4 °C and measured within 24 h after sampling using infrared gas analyses (IRGA, ABB Advance Optima, Frankfurt, Germany; as in van Bergen et al. 2020). Samples for dissolved organic carbon (DOC) were filtered (Ø 0.45 μm) and analyzed using a Shimadzu TOC-L CPH/CPN Analyser (Shimadzu, Kyoto, Japan). TSS concentrations were determined according to standard methods (APHA et al. 2017). Ammonium, nitrite and nitrate assays were performed with technical duplicates in 96-well microtiter plates. Ammonium was measured at 420 nm on a 96□well fluorescence spectrophotometer after reaction with OPA reagent (0.54% (w/v) ortho-phthaldialdehyde, 0.05% (v/v) β-mercaptanol and 10% (v/v) ethanol in 400 mM potassium phosphate buffer (pH 7.3)) as previously described (In’t Zandt et al. 2018). The reaction volumes were adjusted to the 96 well plate: 10 μL of sample and 200 μL of OPA reagent. Nitrite was measured colorimetrically at 540 nm using the Griess assay. Afterwards, an incubation with vanadium chloride at 60°C reduced all nitrate to nitrite and the sample was measured again at 540 nm (García-Robledo et al., 2014).

### 2.3. Pharmaceutical analyses

Samples (2 mL) for measuring pharmaceutical concentrations were taken at 0, 4, 8, 24, 48 and 96 h after the start of the incubation. Samples were centrifuged and the supernatant was stored at −20 °C until chemical analysis. A detailed protocol can be found in the supplementary material (SI1). In summary, solid phase extraction (SPE) was performed using Oasis HLB 3cc SPE cartridges (Waters, Milford, MA, USA) to recover the pharmaceuticals present in the samples. Methanol was used to elute the pharmaceuticals and after evaporation, they were dissolved in 0.1 % v/v formic acid. To extract metformin, a liquid-liquid extraction was performed (Yoshida and Akane 1999) by adding acetonitrile and sodium dodecyl sulphate. Calibration standards (*n* = 6, 0.5-50 nM) were freshly prepared and extracted in the same way as the samples. For the analysis of all the extracts, liquid chromatography tandem mass spectrometry analysis (LC-MS/MS) was used. TargetLynx LC-MS/MS data acquisition software (Waters, Milford, MA, USA) was used for the linear curve fitting. Quadratic curve fitting (y = ax^2^ + bx + c) was applied for the quantification of metformin because standard concentrations of extracted metformin did not fit to a linear model. Optimised LC-MS/MS parameters are provided in the supplementary data (Table S1, S2, and Figure S1)

### 2.4. Biodegradation rate constants

#### 2.4.1 Theory

Biodegradation rate constants are often assumed to follow Michaelis-Menten kinetics (Schwarzenbach et al. 2005), where the reaction rate is a result of enzyme binding, product formation and dissociation of the enzyme-substrate complex (Michaelis and Menten 1913), therefore depending on substrate concentration (Equation 1).

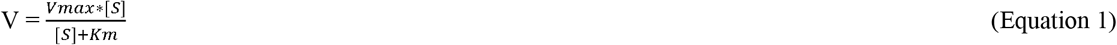

Where V is the reaction rate (mol L^−1^ h^−1^), V_max_ is the maximum reaction rate (mol L^−1^ h^−1^), [S] is the substrate concentration (mol L^−1^) and Km is the substrate concentration were V is half of V_max_ (mol L^−1^). As pharmaceutical concentrations in the wastewater are very low, it can be assumed that the substrate concentration is far below K_m_ and the reaction rate is linearly proportional to the substrate concentration. Hence, biodegradation rate constants can be calculated via pseudo-first order biodegradation kinetics (Schwarzenbach et al. 2005):

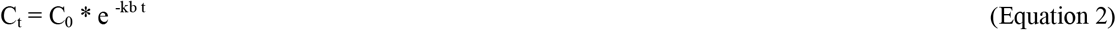

Where C_t_ is the concentration at time t (nM), C_0_ is the concentration at time 0 (nM) and k_b_ is the biodegradation rate constant (h^−1^). Theoretically, the k_b_ (that includes the effect of substrate concentration at each time point) should be constant when the substrate concentration is below K_m_

#### 2.4.2 Biodegradation analyses

The k_b_ was calculated based on the average concentration of three technical replicates over six timepoints (n=6). Exponential decay models were fitted to the concentration over time (Eq. 2) and quality of the model fit was evaluated by visual assessment and by referring to the goodness of fit parameters R^2^ and the p-value. The k_b_ was estimated using R (Version 4.0.2; R Core Team, 2020) and the packages drc, nlme and aomisc (Pinheiro et al. 2017, Ritz et al. 2015). For some pharmaceuticals, only part of the curve was used for model fitting and prior to analyses one outlier was removed from the data (SI 2). Next to first-order biodegradation rate constants, zero order and second order models were fitted as well, but first-order models showed the best overall fit, so we chose to further work with first-order k_b_ values, also for comparability reasons. When we did not observe a significant decrease in concentration, we assumed no biodegradation occurred. Furthermore, first order biodegradation rate constants were correlated with pharmaceutical concentration.

#### 2.4.2 Sorption analyses

To check whether gradual abiotic removal processes (hydrolyses, sorption) occurred in the inactivated sludge treatment, similar first order models were applied for that treatment. When we found a significant rate constant, we subtracted this from the k_b_ of the activated sludge treatment in order to obtain the biodegradation rate constants, thereby excluding chemical processes. Additionally, the solids water partition coefficient (K_d_) for each chemical based on the inactivated sludge treatment were calculated, assuming instantaneous sorption, according to equation 3 (as in Helbling et al. 2012):

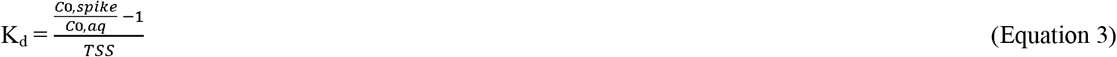

where K_d_ is the solids partitioning coefficient (K_d_, L g^−1^); where TSS is the total suspended solids concentration in the sample g ss L^−1^. The value for C_aq_ was directly measured in the aqueous phase for each sample from the glass bottles (t=0; nM). Assuming instantaneous sorption, K_d_ can be related to the known spike concentration (C_0,spike_; nM) and C_0,aq_. All average values are given with their SD (±1SD).

### 2.5. Molecular analyses

#### 2.5.1. Sampling and DNA isolation

Samples were taken from the inoculum before the experiments (in technical triplicates) and from the bottles at the end of the experiments. Sampling consisted of pipetting 2 mL of activated sludge suspension after thorough mixing. Afterwards, samples were centrifuged for 4 min at 20,000 x *g*: supernatants were transferred to another tube for chemical analysis and pellets were stored at −20□°C until further analysis. DNA was extracted and purified using the DNeasy PowerSoil kit (QIAGEN Benelux) following the manufacturer’s instructions. DNA concentrations were determined using the Qubit dsDNA HS Assay kit and a Qubit fluorometer, both from Thermo Fischer Scientific (Waltham, MA USA).

#### 2.5.2. Bacterial 16S rRNA gene sequencing and analysis

DNA samples were submitted to Macrogen (Seoul, South Korea) for amplicon sequencing of the bacterial 16S rRNA gene hypervariable V3 and V4 region. Sequencing was performed on an Illumina Miseq. The primers used for amplification were Bac341F (5′□CCTACGGGNGGCWGCAG□3′) and Bac785R (5′□ GACTACHVGGGTATCTAATCC □3′) (Klindworth et al. 2013). Subsequent analysis of the sequencing output files was performed with R version 3.4.1 (R Core Team 2020). Pre-processing of the sequencing data was done using the DADA2 pipeline (Callahan et al. 2016). Taxonomic assignment of the reads was done up to the species level when possible using the Silva non-redundant database version 128 (Yilmaz et al. 2014). Count data were normalized to relative abundances. Data visualization and analysis were performed using phyloseq and ggplot packages (McMurdie and Holmes 2013, Wickham and Wickham 2007). All sequencing data were submitted to the GenBank database under the BioProject ID PRJNA641582 (https://www.ncbi.nlm.nih.gov/bioproject/641582) **[available upon acceptance of manuscript; reviewer link: https://dataview.ncbi.nlm.nih.gov/object/PRJNA641582?reviewer=spgi2sb233j9252q74937qeejk]**. Chao1, Simpson and Shannon diversity indices were calculated using the estimate richness function of the phyloseq package. Permutational multivariate analyses of variance (PERMANOVA) was performed using the adonis function of the vegan package (Dixon 2003). The main drivers of differences in the microbial community composition between summer and winter inocula were identified using the top 15 eigenvalues from PERMANOVA. T-tests were performed on individual taxa to check if their changes were significantly different between sample groups.

#### 2.5.3. Quantitative PCR

The relative cq numbers of *amoA* gene/16S rRNA gene copy numbers in inocula and samples from experiment bottles spiked with 30 nM or no pharmaceuticals were obtained using quantitative PCR. The above primers were used for 16S rRNA gene amplification. For bacterial *amoA* gene amplification, the following primers were used: (5’-GGGGTTTCTACTGGTGGT-3’) and (5’-CCCCTCKGSAAAGCCTTCTTC-3’) (Rotthauwe et al. 1997). A detailed qPCR protocol can be found in the supplementary material (SI3).

## 3. Results

### 3.1. WWTP and incubation conditions

Table 2 gives an overview of the characteristics of AS inocula used for the summer and winter experiments. WWTP conditions such as oxygen concentration, DOC and TSS were largely similar in summer and winter at the time of sampling. The water temperature at the WWTP was higher during summer, as well as the DIC concentration, while the concentration of NH ^+^ was higher during winter. During both experiments, samples were incubated at 15 °C, with an oxygen concentration of approximately 1 mg L^−1^ and a constant pH of 7 to prevent ionization changes. During both experiments, TSS slightly increased (3.6-4.1 g ds L^−1^). In general, the background concentration of pharmaceuticals in the AS was similar (Table 3), with exception of the concentration of metformin, which was almost four times higher in winter (40 nM).

**Table 2.** Activated sludge conditions prior to the summer and winter experiment.

|  | Summer | Winter |
| --- | --- | --- |
| Ammonium (μM) | 168 ± 14 | 597 ± 51 |
| Nitrate (μM) | 2 ± 0.6 | 0 |
| Oxygen (mg L<sup>-1</sup>) | 1.3 | 1.5 |
| Temperature (°C) | 18.3 | 13.6 |
| pH | 6.5 | 7 |
| TSS (g ds L<sup>-1</sup>) | 3.9 ± 0.03 | 3.2 ± 0.1 |
| DOC (mg L<sup>-1</sup>) | 45.2 ± 11.4 | 40 ± 7.9 |
| DIC (mg L<sup>-1</sup>) | 19.1 ± 0.8 | 59.7 ± 2.3 |

**Table 3.**
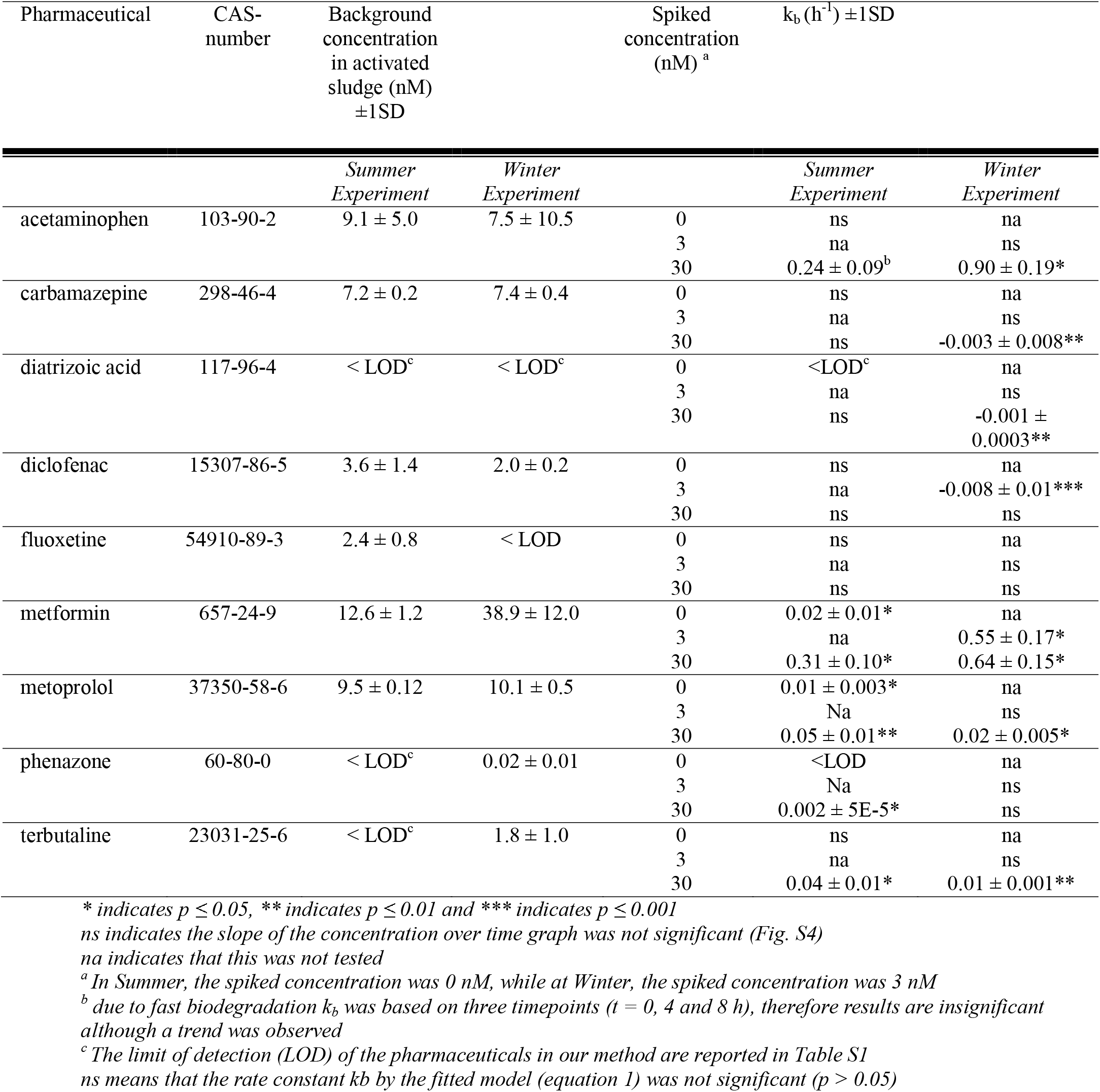
Pharmaceutical background concentrations, k_b_ constants (h^−1^, in case of activated sludge treatments) of summer and winter experiments per treatment. AS is the activated sludge treatment without spiking (summer) or spiked with 3 nM pharmaceuticals (winter) and AS30 is the activated sludge treatment spiked with 30 nM pharmaceuticals. All data points and models used to estimate k_b_ can be found in Figure S4 and details on the statistical analyses can be found in Table S3.

### 3.2. Nitrification activity

As previously mentioned, the ammonium concentration in the winter inoculum was higher than in summer. All ammonium was completely consumed in active bottles (AS and AS30) in both experiments during the first day (Figure S2). Nitrification activity in summer and winter experiments was quantified via nitrate production since ammonium was probably also consumed by heterotrophic bacteria for biomass production. Nitrite was always below the detection limit (2.5 μM) in active bottles. In the summer experiment, the production of 140 μM nitrate was observed in the first two days. In the winter experiment, an increase of 274 μM nitrate was measured mostly during the last three days of the experiment. Bottles with autoclaved biomass did not produce any nitrate (all data shown in Supplementary Figure S3). The nitrification rate was 17 μmoles nitrate day^−1^ g TSS^−1^ in the summer experiment and 28 μmoles nitrate day^−1^ g TSS^−1^ in the winter experiment.

### 3.3. Solid water partition coefficients (Kd)

Solid water partition coefficients were calculated based on instantaneous sorption. We found the highest K_d_ for fluoxetine (0.6 ± 0.1 L g^−1^), followed by carbamazepine (0.03 ± 0.06 L g^−1^), diclofenac (0.02 ± 0.02 L g^−1^) and metoprolol (0.01±0.2 L g^−1^). Results of the autoclaved inactivated sludge treatment showed that most pharmaceutical concentrations did not decrease in this treatment. Therefore, gradual sorption over time was limited and no chemical processes such as hydrolysis were observed. Only a small but significant decrease of diclofenac was observed in the inactivated sludge treatment, likely due to gradual sorption (0.004 ± 0.001 h^−1^, p < 0.01).

### 3.4. Biodegradation rate constants

Table 3 shows background concentrations of pharmaceuticals at the start of the summer and winter experiment and the biodegradation rate constants. No lag phase was observed before biodegradation of pharmaceuticals occurred. Acetaminophen and metformin had a high biodegradability in both experiments. Within 48 hours the concentration of metformin was below the limit of detection (LOD), while for acetaminophen the concentration started to increase again (Figure S4). At the start of the summer experiment (t = 0 h, < 20 minutes after addition of pharmaceuticals), the concentration of acetaminophen was much lower than expected and it was even below LOD in the winter experiment indicating a rapid turnover or uptake by the cells. Biodegradation of acetaminophen was only observed in the treatments with the highest concentration (AS30), while biodegradation of metformin was observed in the low and high concentration treatments (spiked with 3 and 30 nM). The k_b_ of metformin increased at higher concentrations of pharmaceuticals in both experiments. Metoprolol and terbutaline were biodegraded in both experiments at lower rates than acetaminophen and metformin. In the summer experiment, the k_b_ of metoprolol increased at a higher concentration of pharmaceuticals, while at the winter experiment biodegradation was only observed in the high concentration (spiked with 30 nM). Terbutaline was only biodegradable when activated sludge was spiked with a high concentration of pharmaceuticals (30 nM). No biodegradation of fluoxetine was observed. Similarly, phenazone was not biodegraded nor adsorbed during the winter experiment. In the summer experiment, we observed a low biodegradation rate for phenazone (0.002 ± 5E-5 h^−1^) when activated sludge was spiked with 30 nM pharmaceuticals. Diclofenac was not biodegraded in both experiments and when activated sludge was spiked with a low concentration of pharmaceuticals (3 nM), we even observed an increase in the diclofenac concentration.

### 3.5. Relationship of k_b_ with concentration and sludge seasonality

In general, pharmaceuticals with a higher background concentration showed a higher k_b_ value (n=5 pharmaceuticals; difference between pharmaceuticals; Figure 2). Of these compounds, metformin and metoprolol show an increase in k_b_ with concentration (difference within pharmaceuticals). Small effects of seasonality on k_b_ were observed. In the winter time, the k_b_ of acetaminophen was higher than in summer. In winter time, the k_b_ of metformin was higher on average than in summer time, although standard deviation error bars overlap, the 95% CI did not, indicating a significant difference. Metoprolol and terbutaline had only a slightly higher k_b_ in summer than in winter time (note that Figure 2 shows the log y-axe). Phenazone was slightly biodegraded in summer, but not in winter time. Metformin and metoprolol showed a different exponential decrease over time depending on initial concentration in the experiment as a result of spiking and season (Table 3; Figure S4).

**Figure 2.**
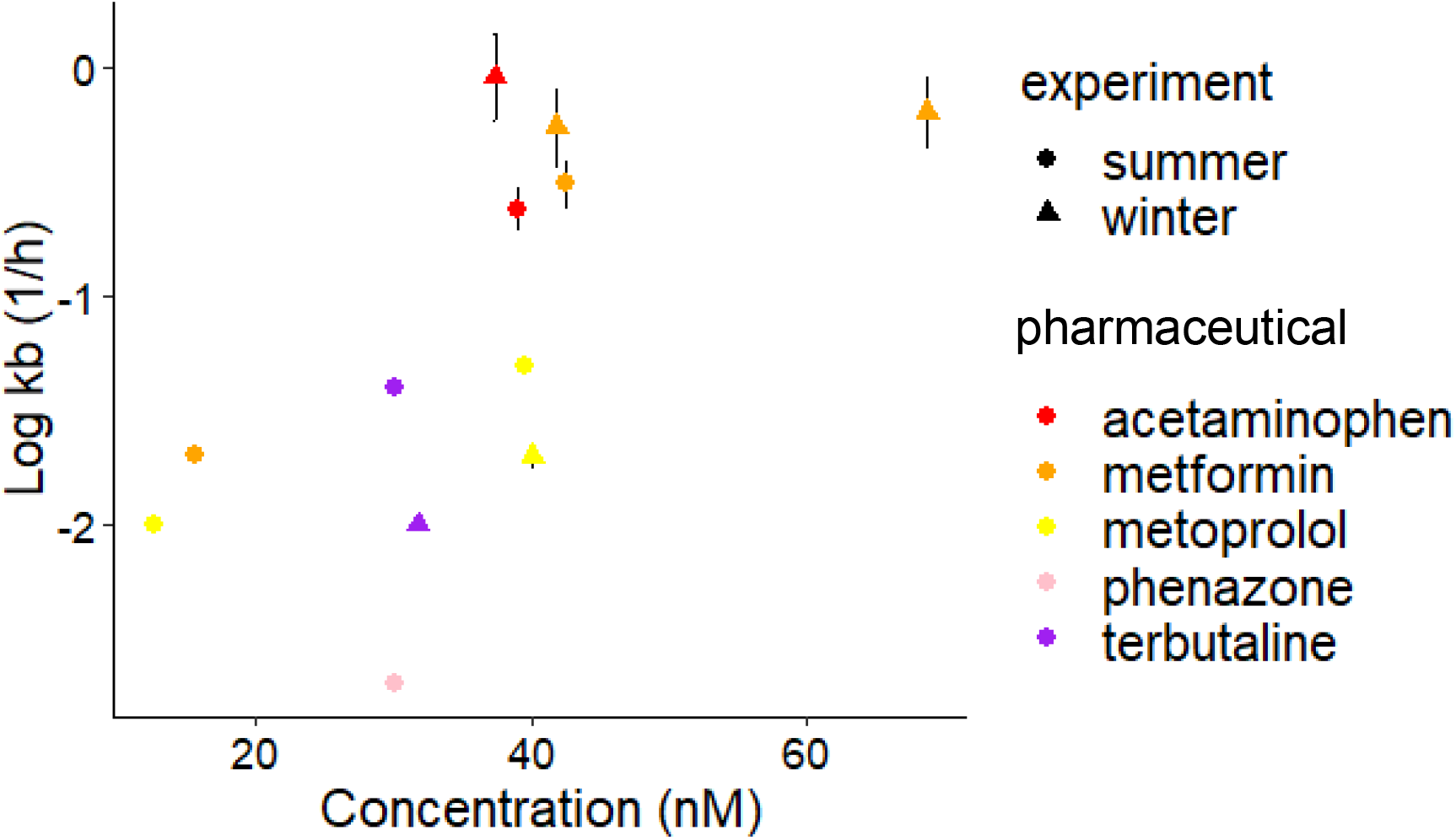
The pseudo-first order biodegradation rate constants k_b_ (log h^−1^) plotted against initial pharmaceutical concentration (nM) at t = 0 of the experiment. All average values are plotted with their SD, but sometimes the SD is too small to see. Colors represent different pharmaceuticals, while circles indicate summer values and triangles indicate winter values.

### 3.6. Microbial community analysis

Microbial community analysis based on bacterial 16S rRNA gene sequencing was first performed on summer and winter inocula. Alpha diversity indices (Chao1, Simpson, Shannon) were not significantly different between summer and winter inocula (p-value>0.05). PERMANOVA testing was used to calculate the significance of compositional differences between the groups. No significant difference was found between inocula (p-value > 0.05 at different taxonomic levels). The bacterial community composition is visualized in a relative abundance bar plot at the lowest taxonomic level (Figure 3). Overall, the data show that the microbial communities of summer and winter inocula closely resemble each other. Zooming into the low differences between inocula (based on PERMANOVA calculated eigenvalues), the phyla contributing the most to variation were *Bacteroidetes*, *Firmicutes*, and *Actinobateria*. All of them were significantly different between both sampling times except for *Bacteroidetes*. At family level, the main contributors were *Burkholderiaceae, Saprospiraceae, Cellvibrionaceae, Haliangiaceae*, and *Rhodanobacteraceae*. Previous studies suggested a putative role of several microorganisms in acetaminophen degradation in AS (Akay and Tezel 2020, Chopra and Kumar 2020, De Gusseme et al. 2011, Palma et al. 2018, Park and Oh 2020a, b, Park and Seungdae 2020, Żur et al. 2018). Among them, the genera *Dokdonella, Flavobacterium, Acinetobacter*, and *Enterococcus* showed a significant (p-value < 0.05) relative abundance increase in winter correlating to a higher biodegradation rate constant of acetaminophen in our experiments (Figure S6 and Figure 2). Microbial community analysis was also performed on biomass at the end of each experiment. The microbial community composition of the inocula and the biomass after 96 h of incubation were still very similar (PERMANOVA p-value>0.05). There were also no significant microbial community changes between experiment bottles with different pharmaceutical concentrations (PERMANOVA p-value>0.05). Differences in bacterial *amo*A gene abundance between inocula and at the end of the bottle incubations were quantified by qPCR and compared as relative cq values using the bacterial 16S rRNA gene cq as internal standard. A small but significant decrease (4%) in the relative cq number of *amo*A/16SrRNA was observed in the winter inoculum (Figure S7). This result aligns with the higher ammonium concentration found in Groesbeek WWTP due to decreased nitrification during winter. As a result of the higher ammonium concentration present at the beginning of the winter experiment, the nitrification rates in the bottles were higher than during the summer experiment.

**Figure 3.**
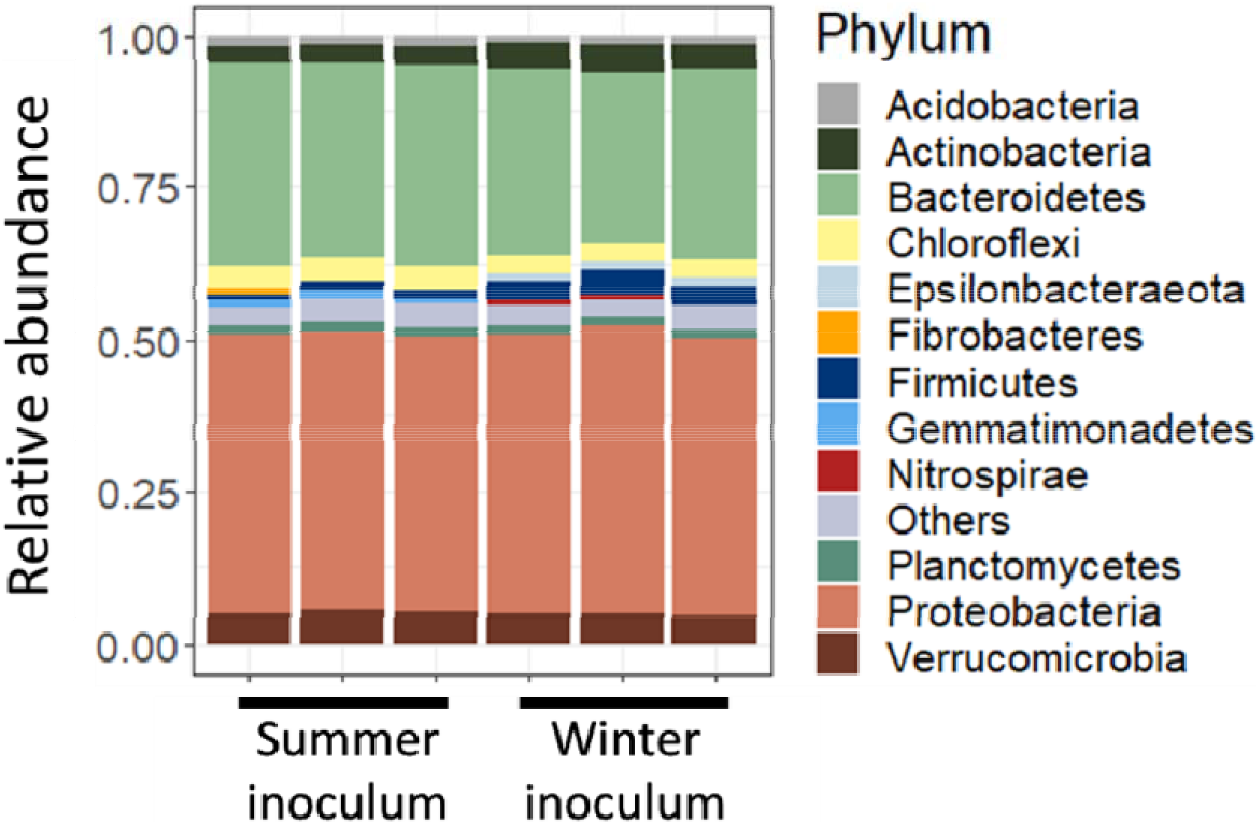
Phylum relative abundance of summer and winter inocula. No significant difference in microbial composition was observed between both inocula (PERMANOVA p-value > 0,05).

## 4. Discussion

In this study we evaluated the effect of initial pharmaceutical concentration and AS seasonality on the biodegradation rate constants (k_b_) of nine highly prescribed pharmaceutical compounds. Acetaminophen and metformin had the highest k_b_ followed by metoprolol, terbutaline and phenazone. Diclofenac, carbamazepine, fluoxetine, and diatrizoic acid were not biodegraded at all. The k_b_ values of biodegradable pharmaceuticals were higher at the highest initial concentration, thereby confirming our hypothesis (i). More specifically, for pharmaceuticals metformin and metoprolol, a higher initial concentration increased their k_b_ (Table 3; Figure 2), even though in the calculation of k_b_ the concentration at each point in time is already taken into account (see 2.4.1). For the other biodegradable compounds, the lowest concentration did not result in any biodegradation, which might indicate a mass transfer constrain as previously demonstrated with the well-known pesticide atrazine (Kundu et al. 2019). In general, biodegradable compounds with a higher background concentration in the AS had the highest k_b_ (Table 3). We observed small differences between the k_b_ obtained with summer and winter AS inocula. The k_b_ of acetaminophen and metformin were higher in the winter experiment, while the k_b_ of terbutaline, metoprolol, and phenazone were slightly lower. The background concentration of the pharmaceuticals was mostly similar in winter and summer, only metformin had a three times higher concentration in winter, explaining the higher k_b_ in winter. Nitrification rates were significantly higher in the winter experiment as a result of the higher ammonium concentration present at the beginning of the experiment. This could not be clearly linked to a difference in k_b_ of specific pharmaceuticals, as acetaminophen was likely not degraded by co-metabolism and differences for metformin were low. No significant differences in microbial community composition were observed between the summer and winter inocula (Figure 3; PERMANOVA p-value > 0.05), opposed to our hypothesis (ii) and not between the start and the end of the experiment. Only the relative abundance of putative acetaminophen degraders (*Dokdonella, Flavobacterium, Acinetobacter*, and *Enterococcus*) was higher in winter – possibly explaining the increased k_b_ of acetaminophen in winter. In general, seasonal effects were not as straightforward as expected in our hypothesis (ii) and our results show pharmaceutical-dependent effects of sludge seasonality in biodegradation rate constants.

Biodegradation was observed as the predominant removal route for most pharmaceuticals in our experiments, except for fluoxetine, diclofenac, carbamazepine and diatrizoic acid. This lack of observed biodegradation was in line with previous studies, where these pharmaceuticals were either very slowly or not biodegraded or showed a high variability in biodegradation depending on the WWTP (Casas et al. 2015, Fernandez-Fontaina et al. 2012, Haiß and Kümmerer 2006, Kruglova et al. 2014, Petrie et al. 2015, Urase and Kikuta 2005). For diatrizoic acid, carbamazepine and diclofenac, we even found increasing concentrations in the winter experiment, corresponding to a small production rate of 0.001-0.003 h^−1^ (Table 3). This could be due to the reverse reaction of metabolites to the parent compounds (Gonzalez-Gil et al. 2019a, Gonzalez-Gil et al. 2019b, He et al. 2019, Tran et al. 2009) or to a desorption process (Gonzalez-Gil et al. 2018). Fluoxetine, diclofenac, carbamazepine and metoprolol were instantaneously adsorbed to the activated sludge in accordance with other studies (Fernandez-Fontaina et al. 2016, Fernandez-Fontaina et al. 2012, Salgado et al. 2012) and corresponding with the highest K_ow_ values (Table 1). Gradual sorption only occurred for diclofenac. Acetaminophen and metformin showed high biodegradation rate constants, similar to other studies (e.g. Petrie et al. 2015, Poursat et al. 2019a), although no biodegradation was observed at the lowest spiked concentrations for acetaminophen. Metoprolol showed varying k_b_ values in other studies (Kasprzyk-Hordern et al. 2008, 2009, Svendsen et al. 2020) and terbutaline was biodegraded well in other studies (Bueno et al. 2012, Choubert et al. 2011). This corresponds to our findings, although no biodegradation was observed at the lowest spiked concentrations for metoprolol in winter and terbutaline in both seasons. Phenazone mostly shows a low k_b_ in other studies, similar to what we found (Casas et al. 2015, Onesios et al. 2009).

When the pharmaceutical concentration increased, k_b_ increased as well for metformin and metoprolol. A possible explanation for this is previous exposure of the microbial inoculum to the compound, which could lead to a higher biodegradation capacity (Poursat et al. 2019b). The biodegradable compounds metformin and metoprolol both occurred in a high background concentration in the activated sludge (Table 3). Similar effects of pharmaceutical concentration on k_b_ have been observed for metoprolol and citalopram in biofilms (Svendsen et al. 2020), with an increase in k_b_ at environmentally relevant concentrations. It is likely that Michealis-Menten theory is not always sufficient to describe biodegradation at low, environmentally relevant concentrations. Based on Michaelis-Menten kinetics, k_b_ should be constant at low concentrations, when the substrate concentration is far below k_m_ (substrate concentration at half the maximum reaction rate, also see theory 2.4.1). We assume this is the case for environmentally relevant concentrations (in the nM range). A possible explanation for this is that Michealis-Menten theory is only developed for one substrate and one enzyme and the enzyme concentration is not considered to increase over time. However, in activated sludge multiple enzymes could be responsible for the biodegrading the same substrate and enzyme concentrations can increase due to microbial growth (Bilal et al. 2019, Tawfik and S 2010). At higher concentrations (above environmentally relevant in the μM range) inhibition processes might start to play a role. For instance, Svendsen et al. (2020) found that the k_b_ of citalopram and metoprolol was decreasing at concentrations higher than environmentally relevant (25-300 μg L^−1^). Svendsen et al. (2020) furthermore found a decrease in k_b_ with increasing concentration for the ibuprofen, sotalol, atenolol, trimethoprim and diclofenac in the concentration range 0.3-300 μg L^−1^. Wei et al. (2019) also found that the k_b_ of four pharmaceuticals (metronidazole, bezafibrate, ibuprofen and sulfamethoxazole) was negatively influenced by concentrations ranging from 0.1 to 3 μM, which are notably higher than the concentrations tested in this study and likely caused inhibition of microbial processes, especially in case for antibiotics.

In a previous experiment, pharmaceutical biotransformation kinetics differed when inocula were taken from different WWTPs (Helbling et al. 2012). In our assays, we observed small k_b_ differences when the inocula came from the same WWTP at different seasons, summer and winter. Acetaminophen and metformin biodegradation rate constants were higher in the winter experiment, wherein nitrification rate was higher as well (Figures S2; S3). However, acetaminophen degradation has been associated to heterotrophic bacteria rather than ammonia oxidizing organisms (De Gusseme et al. 2011). Furthermore, the relative abundance of putative acetaminophen degraders (*Flavobacterium*, *Dokdonella, Acinetobacter, and Enterococcus*) was significantly higher in the winter inoculum (Figure S6) (Akay and Tezel 2020, Chopra and Kumar 2020, Palma et al. 2018, Żur et al. 2018). A bacterium isolated from activated sludge (*Aminobacter* sp. affiliated with *Phyllobacteriaceae*) has been previously correlated with metformin degradation (Poursat et al. 2019a). However, the 16S rRNA gene of *Aminobacter* was absent from our amplicon dataset, indicating that metformin degradation is not limited to *Aminobacter* in activated sludge microbial communities. Other pharmaceuticals such as carbamazepine and diclofenac showed a higher k_b_ with increasing ammonium concentrations in previous 6-day batch experiments inoculated with nitrifying AS (Tran et al. 2009). Likewise, fluoxetine biodegradation has been previously linked to nitrifying activities in a bioreactor of HRT=3.6 days (Fernandez-Fontaina et al. 2012). However, in our assays these three pharmaceuticals did not biotransform at all. Terbutaline, metoprolol, and phenazone biodegradation rates were slightly higher during the summer experiment, wherein nitrification rates were lower. Interestingly, metoprolol biodegradation was inhibited by nitrification in a previous experiment with biomass from constructed wetlands (He et al. 2018). Although there were no significant differences in total microbial community composition between both inocula (Figure 3), there might have been differences in activity of microorganisms able to degrade these pharmaceuticals. Previous studies have reported different microbial composition between seasons (Ju et al. 2014, Liu et al. 2019), but other researchers including us did not observe that (Isazadeh et al. 2016). This shows that microbial seasonal changes might be dependent on each WWTP. In order to identify which exact microorganisms are responsible for all these pharmaceuticals biodegradation under WWTP conditions, further experiments are needed.

## 5. Conclusion

We can conclude that our test shows environmental realism, as the microbial community did not change during the experiment and can therefore be translated to WWTP conditions. Although we found changes in k_b_ of pharmaceuticals, this could not be largely explained by microbial community composition. Likely microbial activity was responsible instead. Seasonality had mostly a minor effect on k_b_ – except for acetaminophen, which was explained by a higher relative abundance of putative acetaminophen degraders. Our test also shows that even similar biodegradation assays with the same inoculum may lead to a different k_b_, depending on pharmaceutical concentration and that Michaelis-Menten theory is not always sufficient to describe kinetics at very low, environmentally relevant concentrations. Therefore, the pharmaceutical concentration should be taken into account when predicting/measuring the k_b_ in order to reliably predict the fate of pharmaceuticals in the WWTP. The concentration dependency of k_b_ may even be used to model the k_b_ more accurately without knowing exact mechanisms. Methods such as transcriptomics may help to model biodegradation more accurately as it gives information on microbial activity.

## 7. Acknowledgements

The authors thank Stefanie Berger for support with the practical work, Pieter Blom for helping with the ammonium and nitrate analyses and Rob Anzion and Maurice van Dael for the LC-MS/MS analyses. The authors also thank Leon Lamers, for making the infrared gas analyses machine available to us, and Sebastian Krosse for help with the TOC analyses. This work was supported by the NWO-domain TTW [grant number 15759], the Netherlands, and the SIAM Gravitation grant funded by NWO [grant number 024.002.002].

## 8. Author contributions

JH, RvZ, AR, MJ and CW initiated the project; JH, RvZ, AR, MJ, CW, TN, TB and ARM contributed to the conceptual framework. TB and ARM conducted the experiments and data analyses. MG and PS developed the LC-MS/MS method. TB and ARM did the majority of manuscript writing and all authors contributed to improved versions of the manuscript.

