## Supplementary information for "Do initial concentration and activated sludge seasonality affect pharmaceutical biodegradation rate constants?"

### **SI 1. Extended methodology pharmaceuticals**

**Chemicals and reagents**

Pharmaceutical standards were purchased in solid form from Sigma Aldrich/Fluka Analytical (Zwijndrecht, the Netherlands) and Merck Group (Darmstadt, Germany). The deuterated internal standards were acquired from Toronto Research Chemicals (North York, Canada). Chemicals needed for LC-MS/MS analysis, formic acid (FA), methanol (MeOH) and acetonitrile (ACN) were purchased at Merck Group (Darmstadt, Germany) and Boom B.V. (Meppel, the Netherlands) respectively. Ammonium formate (NH4HCO2, ≥ 99% HPLC) needed for buffer solution, was retrieved from Aldrich/Fluka Analytical (Zwijndrecht, the Netherlands).

**Pharmaceutical extraction and analysis**

For the analytical determination of micropollutant concentrations in wastewater, an optimized method was developed in collaboration with RadboudUMC. Samples (2.0 mL) were taken at time-points 0, 4, 8, 24, 48 and 96 hours after the start of incubation. Wastewater samples were centrifuged and the supernatant was stored at -20 °C until chemical analysis. To recovery the analytes present in the wastewater, solid phase extraction (SPE) was performed using Oasis HLB 3cc SPE cartridges (sorbent bed of 60 mg from Waters Corporation (Milford, USA)). First an acidic buffer solution with a of pH ~2.2 was prepared, by dissolving 10 mM NH4HCO2, and 3.25 mL FA in 250.0 mL Milli-Q water. Subsequently the SPE cartridges were conditioned with 2.0 mL of MeOH and flushed with 2.0 mL of acidic buffer solution prior to sample loading. SPE extraction was applied on 1.0 mL supernatant, spiked with 0.1 mL of deuterated internal standards (50 µg/L in methanol), diluted in 4.0 mL acidic buffer. Afterwards, 5.0 mL of MeOH was used to elute the retained OMPs. Vacuum was applied on each step. Next, the extracts were evaporated under a moderate stream of nitrogen using a sample concentrator set at 40 oC. Sample extracts were reconstituted in 0.1 % v/v FA (mobile phase A). To extract metformin, a liquid-liquid extraction was performed (as in Yoshida and Akane 1999) (SI 3). 0.1 mL Sodium dodecyl sulfate (2.0 mM) was added to 0.5 mL of wastewater including 1.0 mL ACN. The tubes were placed in the freezer at -20 °C for 30 min and the upper organic layer was taken for analysis. SPE and LLE recovery data was calculated dividing the measured yield concentration by the measured original concentration, see Equation 1

𝑅𝑒𝑐𝑜𝑣𝑒𝑟𝑦 (𝑅%)= (𝐶𝑜𝑛𝑐𝑒𝑛𝑡𝑟𝑎𝑡𝑖𝑜𝑛 (𝑦𝑖𝑒𝑙𝑑)/𝐶𝑜𝑛𝑐𝑒𝑛𝑡𝑟𝑎𝑡𝑖𝑜𝑛 (𝑜𝑟𝑖𝑔𝑖𝑛𝑎𝑙))*100	(Equation S1)

**LC-MS/MS analysis**

For the analysis of pharmaceuticals, Liquid chromatography tandem mass spectrometry analysis (LC-MS/MS) was used. The LC system was coupled to a Xevo TQ-S micro quadrupole mass spectrometer (MS). Electro spray ionization (ESI) was used during the analysis of the pharmaceuticals. LC-MS/MS analysis was operated in positive ion mode to obtain two Multiple reaction monitoring (MRM) transitions per compound. Chromatographic separation was done on an Acquity UPLC BEH C18 (2.1 mm x 100 mm, 1.7 μm) reversed phase column. The mobile phases consisted of eluent (A); 0.1 % v/v FA in Milli-Q water and eluent (B); 100% ACN. Injection volume was set on 2.5 µL, and the gradient program used a flow of 0.5 mL/min. The gradient program was started at 100% (A) for only 0.2 min, changing to 100% (B) at 2.0 min. The 100% (B) was continued till 4.3 min and then switched back 100% (A) at 4.5 min. The overall gradient program was applied for 8 min. Prior LC-MS/MS analysis, calibration standards (n = 6, 0.5 – 100.0 µg/L) were freshly prepared and extracted in the same way as the wastewater samples. TargetLynx LC-MS/MS data acquisition software (Waters Corporation Milford, USA) was used for the linear curve fitting. Furthermore, 1/X weighting was used to minimise the mathematical error for the for the lowest calibration points (Almeida et al. 2002). This weighing was selected because it gave the lowest standard errors for all the points in the calibration curve compared to 1/X^2^ or not weighing. Quadratic curve fitting with the same 1/X weighting (y = ax2 + bx + c) was applied for the quantification of metformin. According to (Liu et al. 2019), the nonlinearity of metformin calibration curve is produced by an interplay between the deuterated internal standard and the original compound. Optimised LC-MS/MS parameters are provided in Figure S1, Table S1, and S2.

### **SI 2. Biodegradation rate constants**

Metformin and acetaminophen where either degraded < 48 h or after that time other processes may have started to play a role, for instance release by microorganisms or back-transformation (Gulde et al. 2016, Tran et al. 2018). Therefore, only part of the curve was used to calculate k_b_ values. In order to estimate k_b_ values for acetaminophen with an exponential decay models (according to Equation 1), the first timepoint was replaced by the theoretical concentration of the spiked treatment (i.e. background concentration + 3 or 30 nM). This was because the first measurement of acetaminophen (t = 0 h, < 20 min after addition) in the spiked activated sludge treatments were much lower than the concentration in the inactivated sludge treatment or even < LOD, likely due to uptake processes. One outlier was removed in order to calculate the k_b_ for metformin in the winter experiment: a value of > 300 nM metformin in the AS3 treatment at time = 8 h, which was over a hundred times higher than the theoretically added concentration (3 nM), matching the other two technical replicates. This value was also outside the interquartile range and this was likely caused by technical sampling error.

### **SI 3. Quantitative PCR protocol**

All qPCR reactions were performed using PerfeCTA Quanta master mix (Quanta Bio, Beverly, MA) and 96‐well optical PCR plates (Bio‐Rad Laboratories Veenendaal, the Netherlands) with optical adhesive covers (Applied Biosystems, Foster City, CA). All reactions were performed on a C1000 Touch thermal cycler equipped with a CFX96 Touch™ Real‐Time PCR Detection System (Bio‐Rad Laboratories, Veenendaal, the Netherlands). The qPCR total volume was 25 µL containing 1 µL DNA sample of 1 to 10 ng/µL, 0.5 µL of each primer solution of 20 µM, 12.5 µL of Quanta master mix and 10.5 µL of autoclaved MiliQ water. Negative controls were added to each run by replacing the template with sterile Milli‐Q water. All qPCR data was analyzed using the Bio‐Rad CFX Manager version 3.0 (Bio‐Rad Laboratories, Veenendaal, the Netherlands). The qPCR protocol consisted of initial denaturation at 95^o^C 3 min followed by 39 cycles of denaturation, annealing, and extending (95^o^C 30 sec, 60^o^C 30 sec, and 72^o^C 30 sec, respectively), and two final steps at 65^o^C 5 sec and 95^o^C 50 sec.

### **Figure S1. Recovery of pharmaceuticals spiked in Milli-Q and synthetic sewage water**

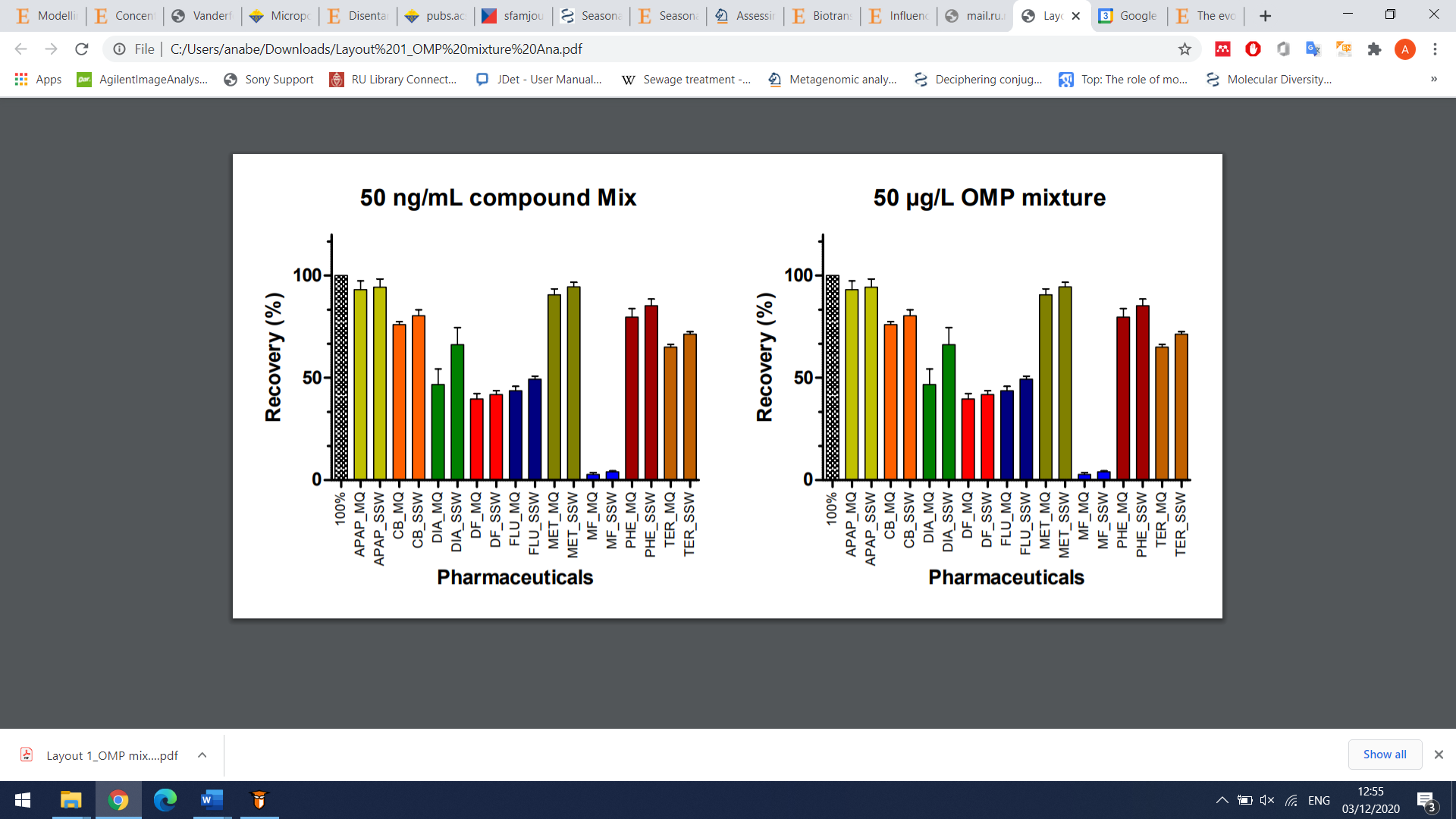

**Figure S1.** Recovery of pharmaceuticals spiked in Milli-Q and synthetic sewage water. A known concentration of 50 µg/L known concentrations, for the pharmaceuticals, acetaminophen (APAP), carbamazepine (CB), diatrizoate (DIA), diclofenac (DF), Fluoxetine (FLU), metoprolol (MET), metformin (MF), phenazone (PHE) and terbutaline (TER).

### **Figure S2. Ammonium profiles during summer and winter experiments**

*Orange colour represents the experiment bottles with 30nM of each pharmaceutical added. Grey colour represents the bottles with doubled autoclaved biomass and 30nM of each pharmaceutical added. Blue colour in summer experiment represents the bottles with no pharmaceuticals added. Blue colour in winter experiment represents the bottles with 3 nM of each pharmaceutical added.*

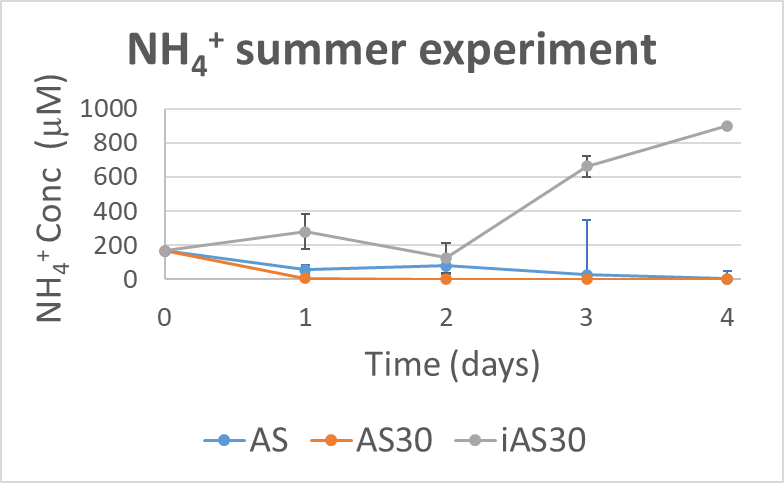

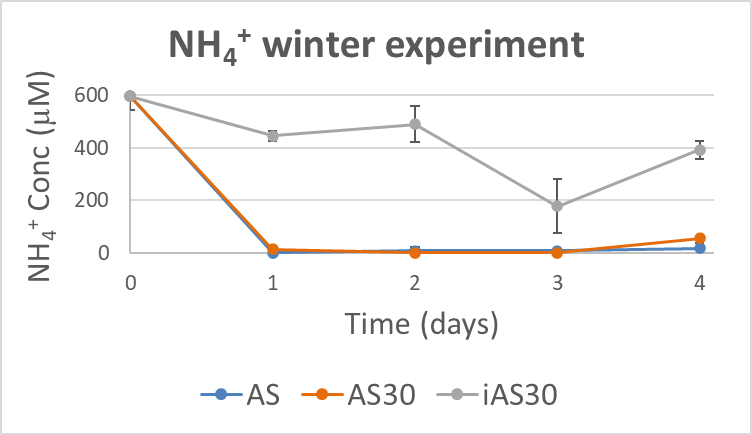

### **Figure S3. Nitrate profiles and production rates during summer and winter experiments**

*Nitrate production rates are calculated by fitting the dots to a line. In the summer experiment the rate is 67.448 µM*day^-1^ with a residual standard error = 53.919 on 13 degrees of freedom, p = 0.0097. In the winter experiment the rate is 90.265 µM*day^-1^ with a residual standard error = 25.4132 on 13 degrees of freedom, p = 2.63E-13.*

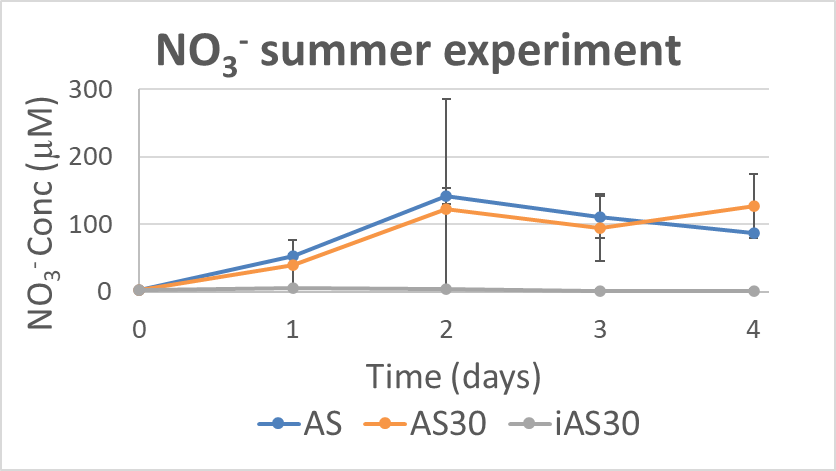

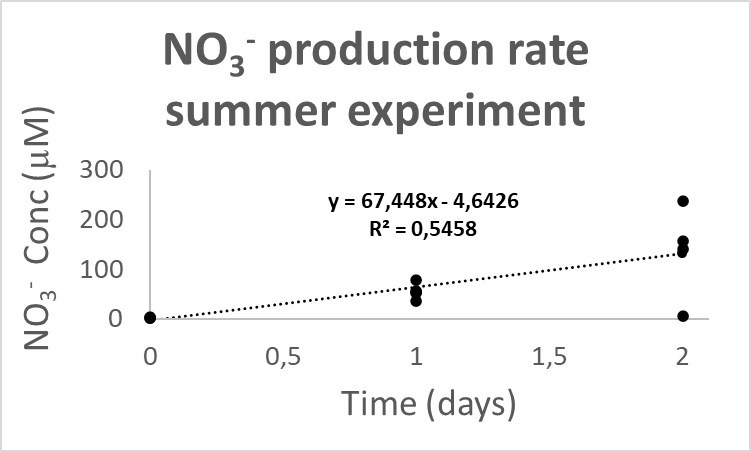

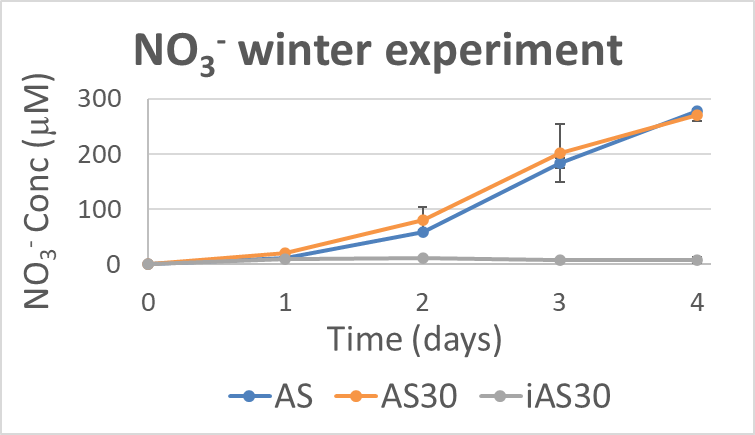

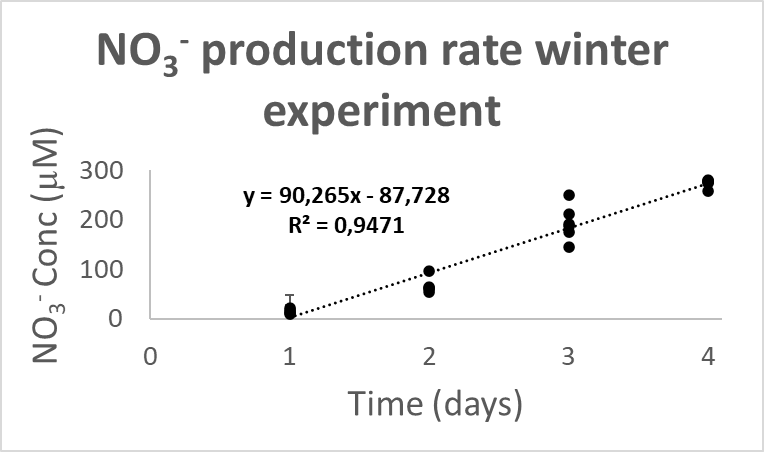

### **Figure S4. Concentrations pharmaceuticals over time during the summer (a) and winter experiment (b).**

Graphs are shown for pharmaceuticals: acetaminophen, carbamazepine, diatrizoic acid, diclofenac, fluoxetine, metoprolol, metformin, terbutaline and phenazone. All measurement points are shown of three treatments: 1) activated sludge with no (summer; AS) or spiked with 3 nM of each pharmaceutical (winter; AS); 2) activated sludge spiked with 30 nM of each pharmaceutical (AS30); and 3) inactivated sludge spiked with 30 nM of each pharmaceutical (iAS30). In some cases, no points are shown in case measurements were < LOD. Significant models are plotted in the graphs.

a) Concentration that was measured over time in the summer experiment
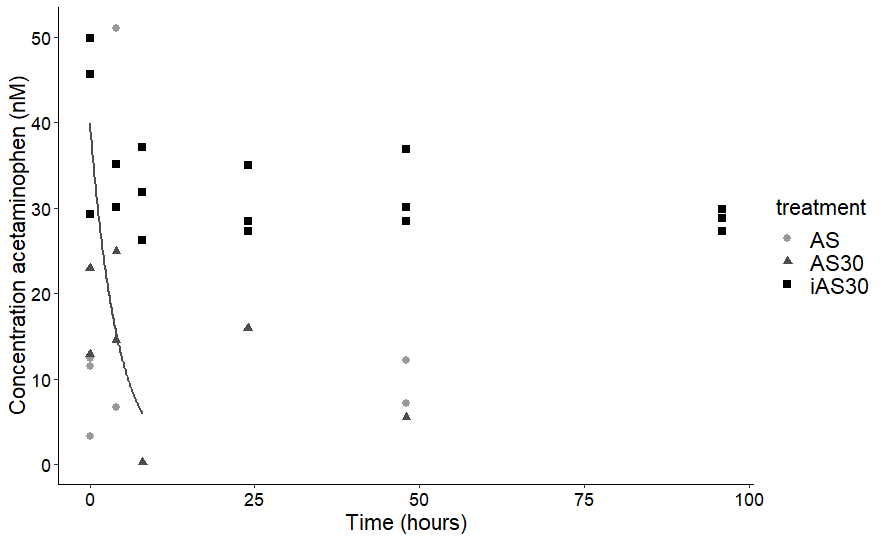

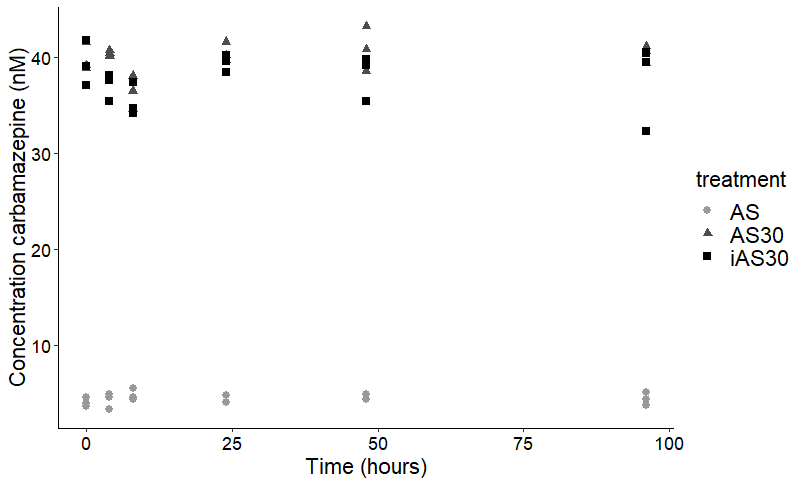

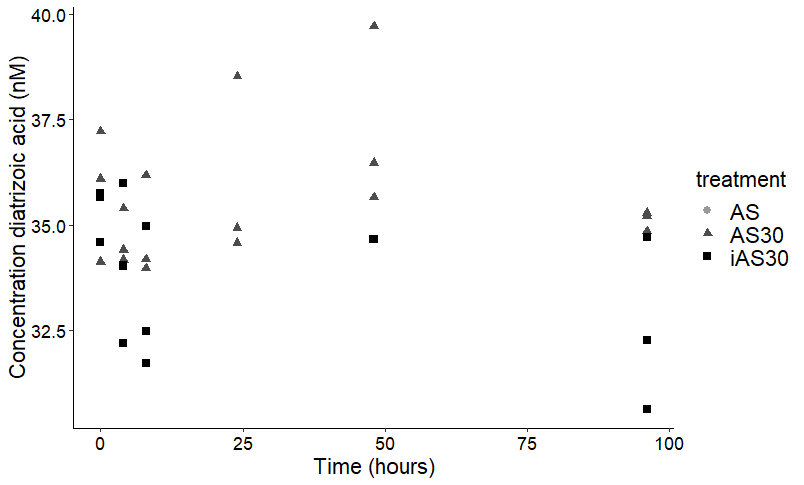

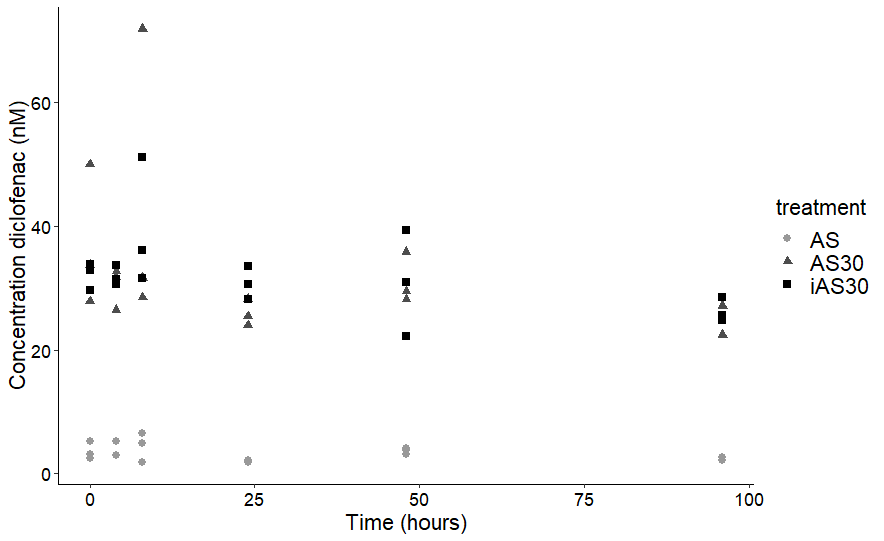

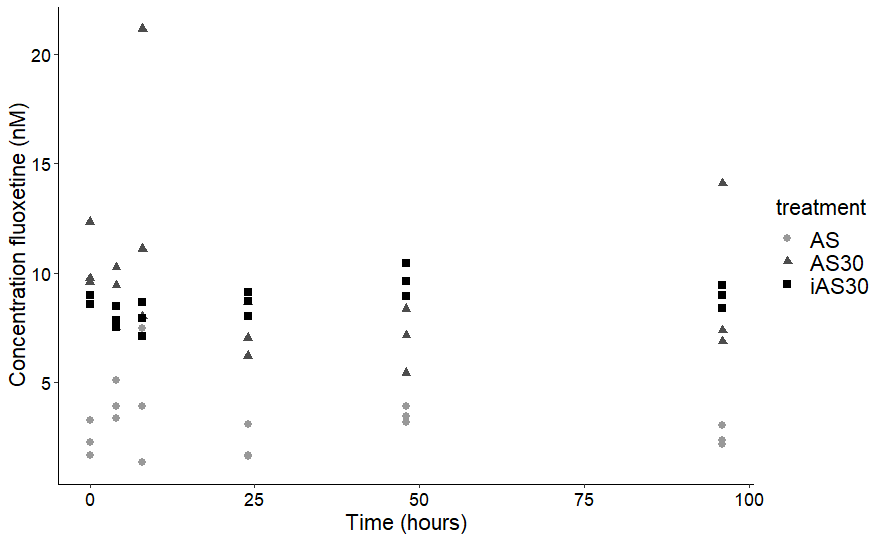

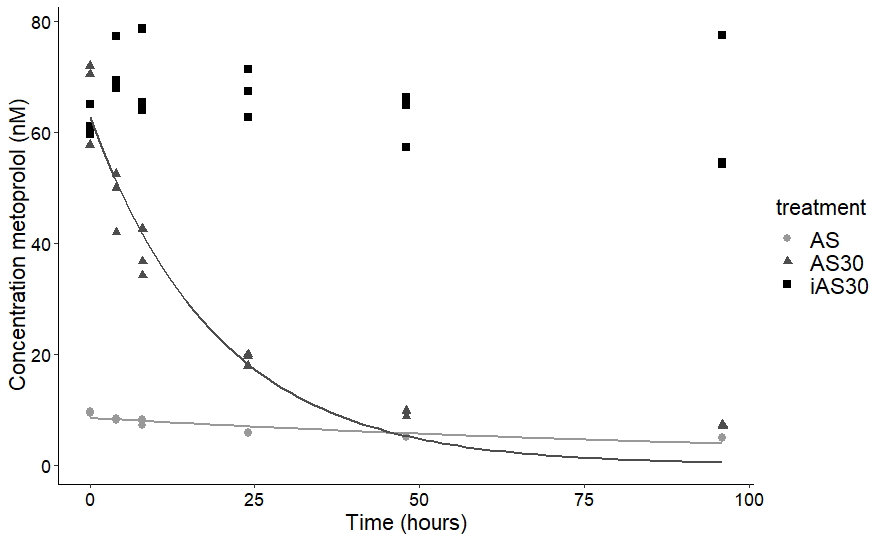

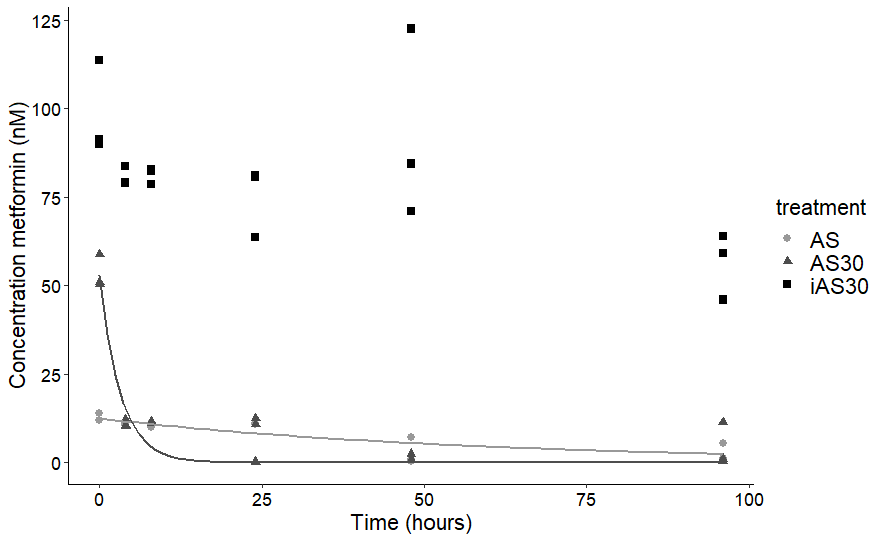

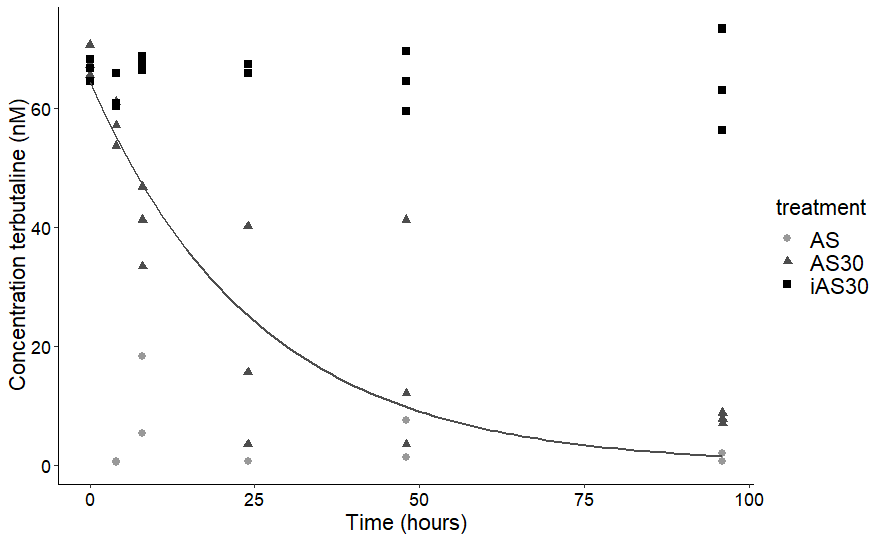

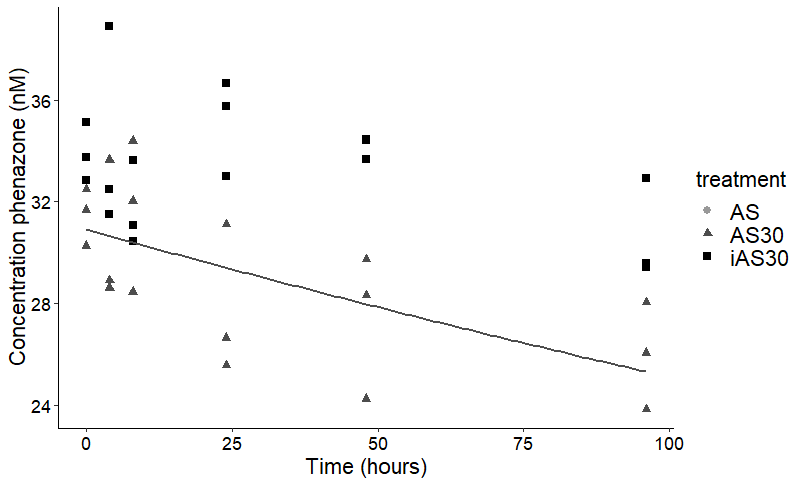

b) Concentration that was measured over time in the winter experiment

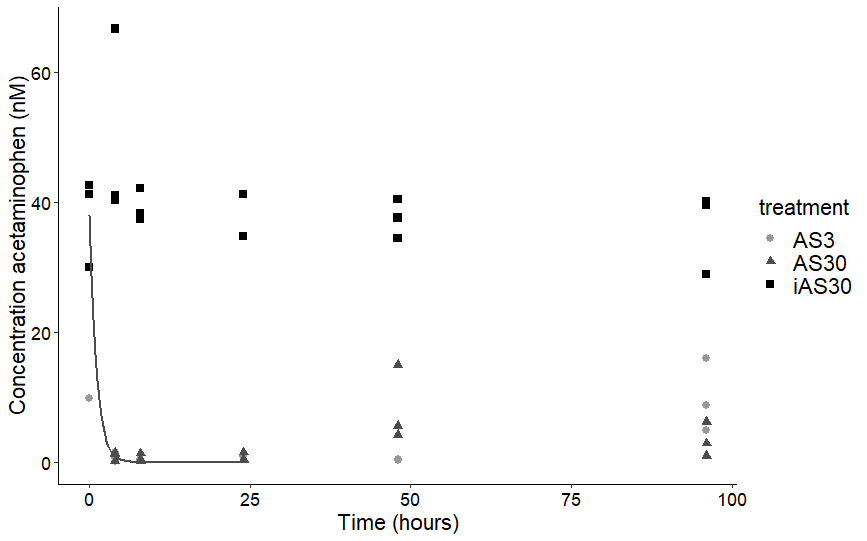

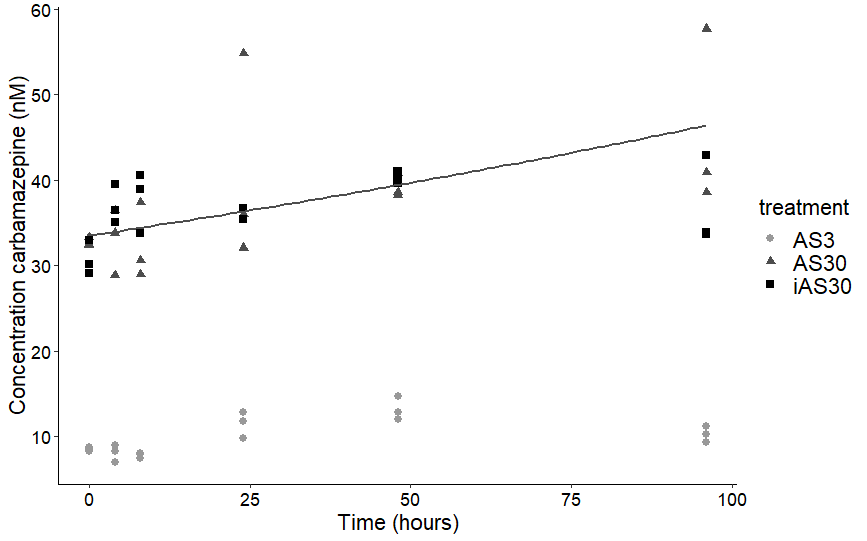

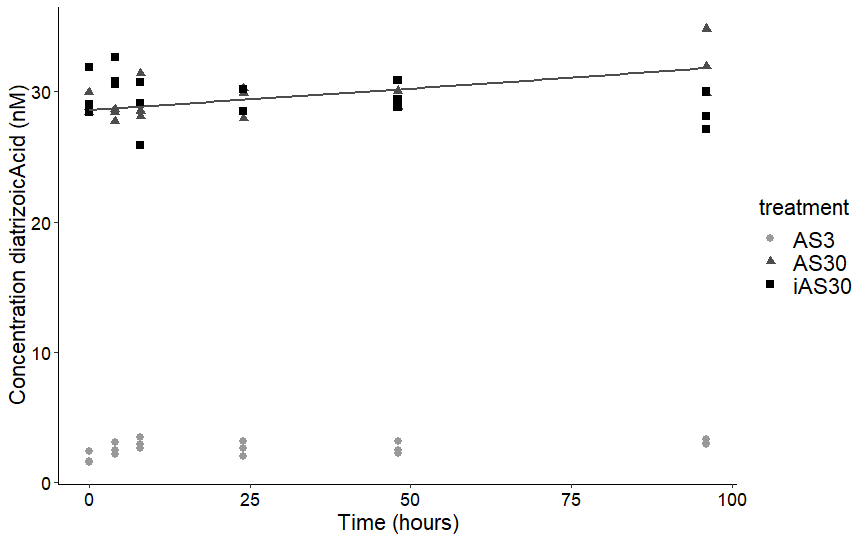

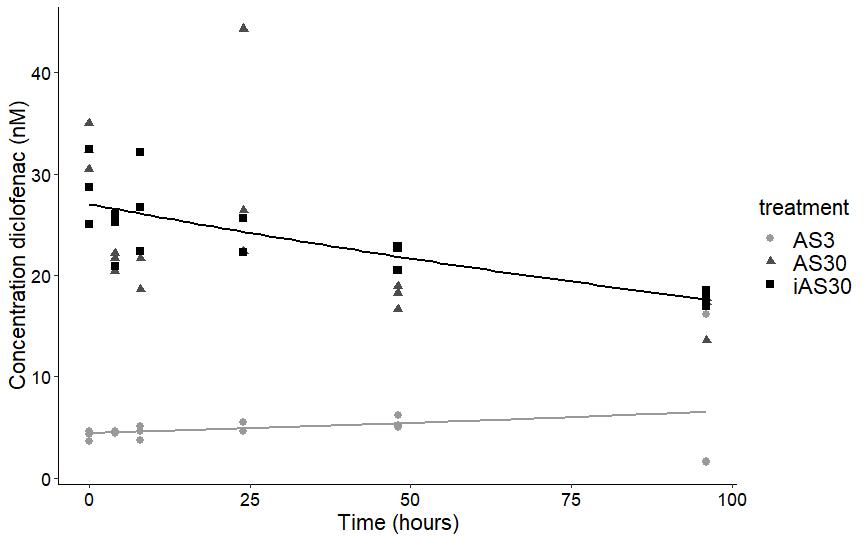

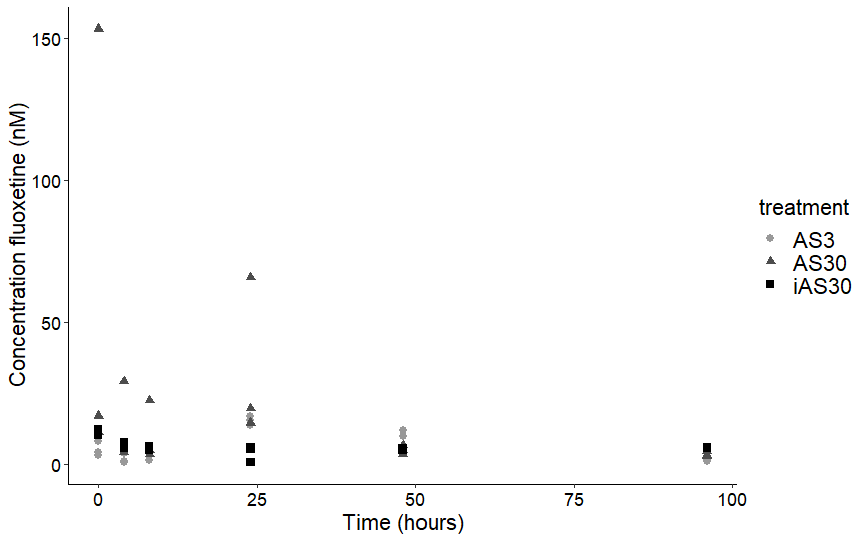

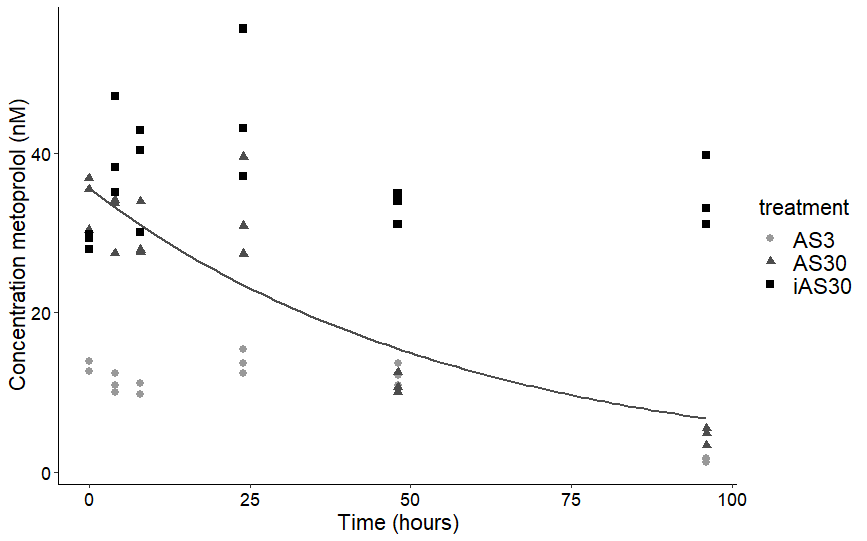

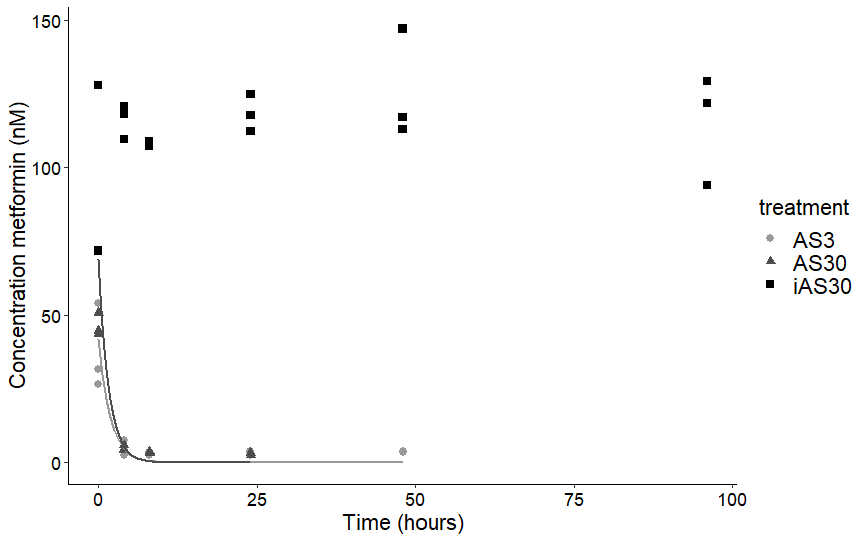

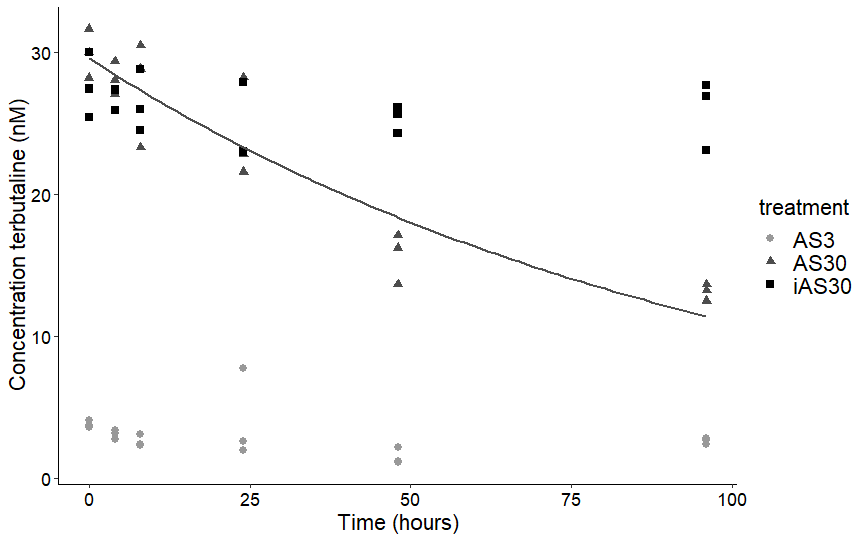

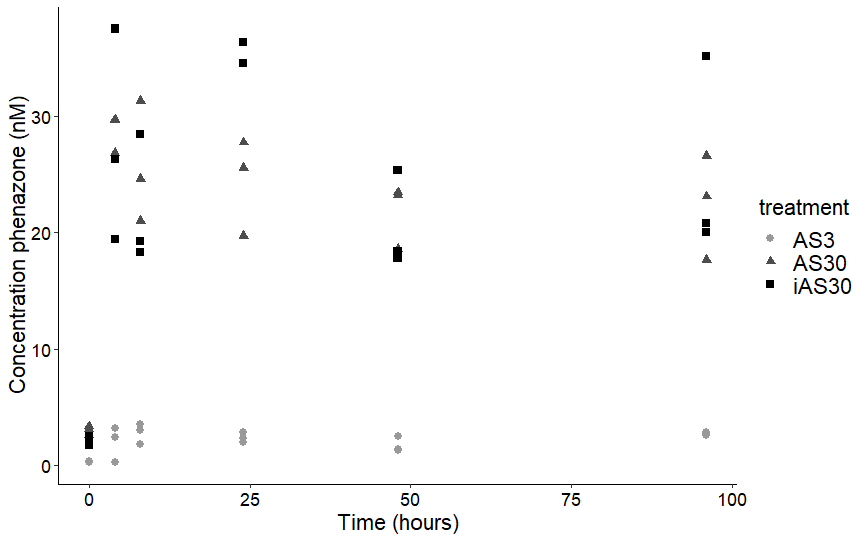

### **Figure S6. 16S rRNA gene relative abundance of putative acetaminophen degraders in summer and winter inocula**

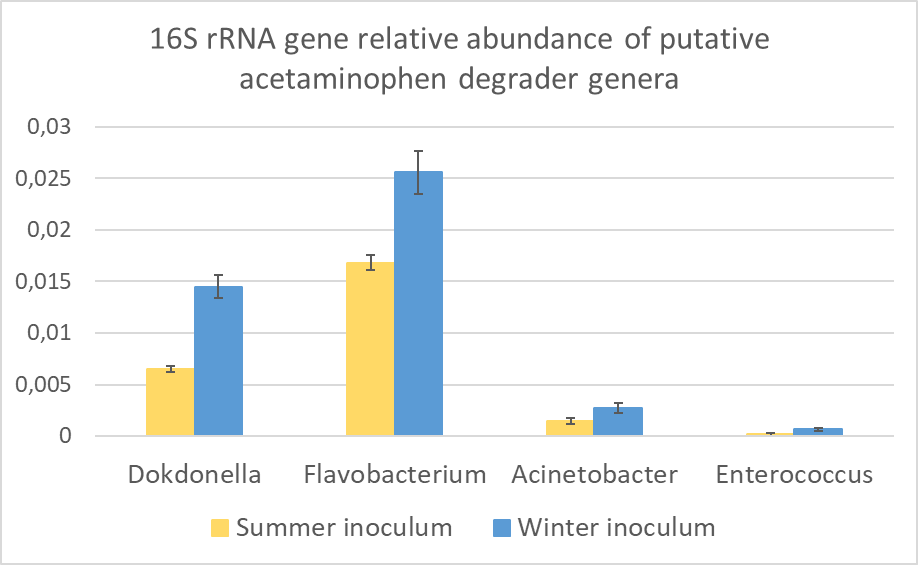

### **Figure S7. Relative cq value of bacterial *amo*A and 16S rRNA genes in summer and winter inocula and experiment bottles with different pharmaceutical concentrations.**

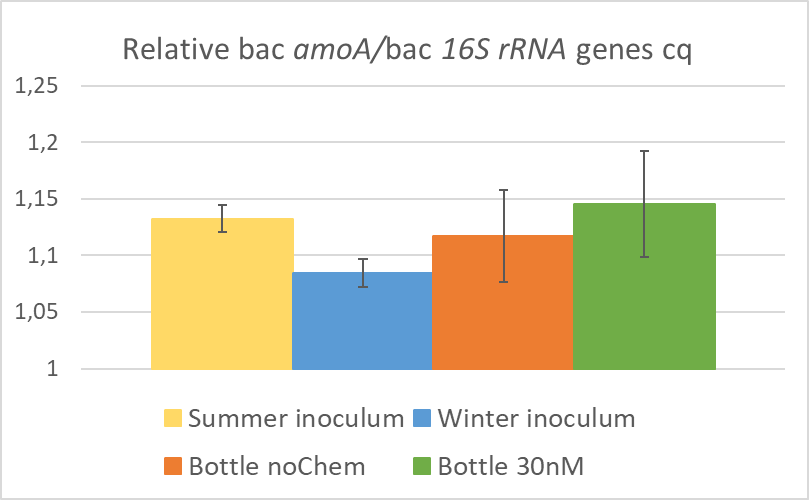

### **Table S1. Optimization parameters LC-MS/MS and SPE extraction**

Optimization parameters LC-MS/MS and SPE extraction

| **Compound** | **Retention**  Time *t_r_*  (min) | **Transition**  [M+H]  (MRM) | **LOD^A^**  (µg/L) | **Milli-Q Matrix**  x̄ SPE Recovery (%) (±sd)  *n* = 3 | **Synthetic Sewage Water**  x̄ SPE Recovery (%) (±sd)  *n* = 3 |
| --- | --- | --- | --- | --- | --- |
| Acetaminophen | 1.69 | (151.968> 109.915)  (151.968> 64.844) | 0.24 | 93.2  (±7.3) | 94.4  (±6.8) |
| Acetaminophen-d4 | 1.70 | (156 > 69) | - | - | - |
| Carbamazepine | 2.26 | (236.968 > 178.876)  (236.968 > 165.019) | <0.01 | 76.1  (±2.4) | 80.2  (±5.8) |
| Carbamazepine-d10 | 2.27 | (247 > 204) | - | - | - |
| Diatrizoate | 1.63 | (614.777 > 360.917)  (614.777 > 233.018) | <0.01 | 46.8  (±13.2) | 66.3  (±14.5) |
| Diatrizoate-d6 | 1.63 | (621 > 367) | - | - | - |
| Diclofenac | 2.66 | (295.904 > 214.804)  (295.904 > 150.951) | 0.09 | 39.6  (±4.6) | 41.9  (±3.0) |
| Diclofenac-d4 | 2.66 | (299.9 > 155) | - | - | - |
| Fluoxetine | 2.15 | (310.224 > 148.035)  (310.224 > 43.848) | 0.03 | 43.7  (±3.8) | 49.3  (±2.7) |
| Fluoxetine-d5 | 2.16 | (315 > 153) | - |  |  |
| Metoprolol | 1.87 | (268.266 > 116.031)  (268.266 > 98.017) | <0.01 | 90.6  (±5.0) | 94.5  (±3.8) |
| Metoprolol-d7 | 1.88 | (275 > 123) | - | - | - |
| Metformin^B^ | 0.57 | (129.968> 70.94)  (129.968> 59.947) | 0.11 | 1.3  (±0.7) | 0.7  (±0.2) |
| Metformin-d6 | 0.56 | (136 > 77) | - | - | - |
| Phenazone | 1.94 | (189.032 > 131)  (189.032 > 103.878) | 0.13 | 79.7  (±7.4) | 85.4  (±4.9) |
| Phenazone-d3 | 1.95 | (192 > 77) | - | - | - |
| Terbutaline | 1.60 | (226.224 > 170.019)  (226.224 > 106.999) | <0.01 | 65.1  (±2.2) | 85.4  (±4.9) |
| Terbutaline-d9 | 1.61 | (235 > 153) | - | - | - |

**^A^** **LOD** = Lowest calibration point detected * (3/ (*s/n*)), determined in Milli-Q 0.1% FA.

**^B^**: Quadratic Curve fitting used; y = ax^2^+bx+c

### **Table S2. LLE extraction for metformin**

LLE extraction for metformin

| **LLE**  **Metformin** | **Milli-Q Matrix**  **x̄ (R%) (±sd)** | **Synthetic Sewage Water**  **x̄ (R%) (±sd)** |
| --- | --- | --- |
| 50.0 µg/L (*n* = 3) | 82.4 (±8.4) | 64.9 (±4.7) |

### **Table S3. Statistics of fitted biodegradation rate constants.**

### K_b_ is the pseudo-first order biodegradation rate constant and DF is degrees of freedom. AS is the activated sludge treatment without spiking (summer) or spiked with 3 nM of each pharmaceutical (winter), AS30 is the activated sludge treatment spiked with 30 nM of each pharmaceutical and iAS30 is the inactivated sludge treatment spiked with 30 nM of each pharamceutical.

|  |  | Summer | Exp. |  |  |  |  |  | Winter | Exp. |  |  |  |  |  | |
| --- | --- | --- | --- | --- | --- | --- | --- | --- | --- | --- | --- | --- | --- | --- | --- | --- |
| OMP | treatment | kb | SD | intercept | SD | p slope | residual standard error | DF | kb | SD | intercept | SD | p slope | residual standard error | | DF |
| Acetaminophen | AS | ns | ns | ns | ns | ns | ns | ns | ns | ns | ns | ns | ns | ns | ns | |
|  | AS30 | 0.24 | 0.09 | 39.94 | 7.1 | 0.23 | 7.23 | 1 | 0.9 | 0.19 | 37.98 | 0.76 | 0.041 | 0.76 | 2 | |
|  | iAS30 | ns | ns | ns | ns | ns | ns | ns | ns | ns | ns | ns | ns | ns | ns | |
| Carbamazepine | AS | ns | ns | ns | ns | ns | ns | ns | ns | ns | ns | ns | ns | ns | ns | |
|  | AS30 | ns | ns | ns | ns | ns | ns | ns | -0.003 | 0.008 | 33.51 | 1.37 | 0.01 | 2.62 | 4 | |
|  | iAS30 | ns | ns | ns | ns | ns | ns | ns | ns | ns | ns | ns | ns | ns | ns | |
| Diatrizoate | AS | ns | ns | ns | ns | ns | ns | ns | ns | ns | ns | ns | ns | ns | ns | |
|  | AS30 | ns | ns | ns | ns | ns | ns | ns | -0.001 |  | 28.6 | 0.35 | 0.01 | 0.64 | 4 | |
|  | iAS30 | ns | ns | ns | ns | ns | ns | ns | ns | ns | ns | ns | ns | ns | ns | |
| Diclofenac | AS | ns | ns | ns | ns | ns | ns | ns | -0.004 | 0.005 | 4.41 | 0.11 | 0.001 | 0.22 | 4 | |
|  | AS30 | ns | ns | ns | ns | ns | ns | ns | ns | ns | ns | ns | ns | ns | ns | |
|  | iAS30 | ns | ns | ns | ns | ns | ns | ns | 0.004 | 0.001 | 27.03 | 0.92 | 0.01 | 1.6 | 4 | |
| Fluoxetine | AS | ns | ns | ns | ns | ns | ns | ns | ns | ns | ns | ns | ns | ns | ns | |
|  | AS30 | ns | ns | ns | ns | ns | ns | ns | ns | ns | ns | ns | ns | ns | ns | |
|  | iAS30 | ns | ns | ns | ns | ns | ns | ns | ns | ns | ns | ns | ns | ns | ns | |
| Metformin | AS | 0.017 | 0.006 | 12.47 | 1.3 | 0.04 | 1.98 | 4 | 0.547 | 0.17 | 41.87 | 3.14 | 0.052 | 3.14 | 3 | |
|  | AS30 | 0.31 | 0.1 | 52.77 | 6.15 | 0.04 | 6.09 | 4 | 0.639 | 0.15 | 68.88 | 3.09 | 0.051 | 3.09 | 2 | |
|  | iAS30 | ns | ns | ns | ns | ns | ns | ns | ns | ns | ns | ns | ns | ns | ns | |
| Metoprolol | AS | 0.008 | 0.003 | 8.48 | 0.59 | 0.03 | 0.96 | 4 | ns | ns | ns | ns | ns | ns | ns | |
|  | AS30 | 0.05 | 0.01 | 62.64 | 4.34 | 0.01 | 5.01 | 4 | 0.017 | 0.005 | 35.55 | 3.41 | 0.03 | 5.304 | 4 | |
|  | iAS30 | ns | ns | ns | ns | ns | ns | ns | ns | ns | ns | ns | ns | ns | ns | |
| Terbutaline | AS | ns | ns | ns | ns | ns | ns | ns | ns | ns | ns | ns | ns | ns | ns | |
|  | AS30 | 0.039 | 0.01 | 64.37 | 6.08 | 0.03 | 7.3 | 4 | 0.01 | 0.001 | 29.57 | 1.04 | 0.002 | 1.68 | 4 | |
|  | iAS30 | ns | ns | ns | ns | ns | ns | ns | ns | ns | ns | ns | ns | ns | ns | |
| Phenazone | AS | ns | ns | ns | ns | ns | ns | ns | ns | ns | ns | ns | ns | ns | ns | |
|  | AS30 | 0.002 | 0.00005 | 30.91 | 0.65 | 0.02 | 1.15 | 4 | ns | ns | ns | ns | ns | ns | ns | |
|  | iAS30 | ns | ns | ns | ns | ns | ns | ns | ns | ns | ns | ns | ns | ns | ns | |
